# Metabolic growth mechanisms and theoretical growth potential of global woody plant communities

**DOI:** 10.1101/2024.10.02.616230

**Authors:** Shumiao Shu, Xiaolu Tang, George Kontsevich, Xiaodan Wang, Wanze Zhu, Yangyi Zhao, Wenzhi Wang, Xiaoxiang Zhao, Zhaoyong Hu

## Abstract

Predicting the growth and maximum biomass (*M*_max_) of woody plant communities (WPC) is challenging due to the complexity and variability of tree growth. While Metabolic Scaling Theory (MST) offers a promising concept, its current theoretical framework is still insufficient. Here, we applied MST principles and our previous findings to propose an iterative growth model for forest growth (IGMF). This model and its extension show that WPC growth, net primary productivity and other carbon budgets - such as total primary productivity, autotrophic respiration, organ turnover biomass and non-structural carbohydrates - can be expressed as functions of current biomass, maintenance respiration rate per unit biomass and stand age or *M*_max_. These functions are globally convergent, allowing us to estimate the current (2018-2020) global *M*_max_ at 1451 ± 26 Pg based on the current state of WPCs alone, with a growth potential of 518 Pg, 83% of which is attributable to shrublands. By the end of the century, climate change is projected to reduce the total *M*_max_ by 266 Pg, mainly in species-rich evergreen broadleaf forests. Further analysis indicates that species richness increases the climate sensitivity of *M*_max_, while soil organic and moisture affects the direction of this response. Our findings reveal WPC growth kinetics and show a shift in the main contributor to terrestrial carbon sequestration from forests to shrublands.

**Significance Statement:** This study introduces a new theoretical model for understanding and predicting the growth and carbon budgets of woody plant communities (WPCs), which applies to diverse WPCs globally and reveals their convergent metabolic growth patterns.

We predict future changes in the maximum biomass of woody plant communities and find a significant decline in evergreen broadleaf forests, where sensitivity and response to climate change are influenced by current species richness and soil conditions.

---

Woody plant communities (WPCs) are important components of terrestrial ecosystems. Their growth is the main driver of terrestrial carbon sequestration. However, consistent understanding of their growth patterns is lacking. A key reason for this is the considerable uncertainty in the growth of woody plants, especially trees. For example, while tree mortality increases with increasing temperature, changes in tree growth are often inconsistent (Chen *et al*. 2016). Without a comprehensive understanding of WPC growth, relying solely on extrapolations from individual growth observations can lead to divergent projections of WPC carbon sequestration (Keenan *et al*. 2014; Piao *et al*. 2014; Charney *et al*. 2016).

A recent study provided a breakthrough in understanding the growth mechanisms of WPCs, showing that their net primary productivity (NPP) can be expressed as a general scaling function of age and biomass (Michaletz *et al*. 2014; Michaletz *et al*. 2018). Although this convergence is based on metabolic scaling theory (MST), the full underlying theory remains unexplored. In particular, it is necessary to elucidate how these variables drive WPC growth and to clarify the relationships between metabolism, gross primary productivity (GPP), respiration, organ turnover, non-structural carbohydrates (NSCs) and mortality. These relationships are critical for assessing and predicting changes in WPC carbon sequestration. We aimed to develop a comprehensive metabolic growth theory for WPCs. This theory would also predict their maximum biomass based on current carbon budgets.

Our recently proposed generalised metabolic growth model for describing individual growth, the iterative growth model (IGM) (Shu *et al*. 2021)(Box.1), offers a promising basis for achieving these goals. Like other metabolic growth models, such as the ontogenetic growth model (OGM), the IGM is also derived from the MST framework and the growth-maintenance respiration paradigm, following two main steps: (1) assuming the total metabolic rate of an organism is approximately proportional to the *b* power of its size (where *b* = 0.75 in the MST); (2) obtaining the growth metabolic rate and growth rate by subtracting the maintenance metabolic rate, which is proportional to the organism’s size, from the total metabolic rate. The key difference is that the IGM takes into account the genetic and physiological processes that inherently control growth and introduces the concept of unit tissue "formation time" (*T*) to reflect the duration of this control. This enhancement reveals the constraints of the second law of thermodynamics on organism growth and the discrete nature of growth (Shu *et al*. 2021) thus obtaining a more complete metabolic growth theory (Box.1). Mathematically, the IGM emphasises that the change in size over time can vary within the limits of the Gonpertz and Richard curves. In contrast, the OGM is only a special case of the IGM (where *b* = 0.75 and *T* approaches 0) and represents the theoretical lower limit of the growth trajectory (Shu *et al*. 2021; Wang *et al*. 2023; Yao *et al*. 2023). Preliminary evidence suggests that the IGM may better explain plant or tree growth than the OGM (Shu *et al*. 2021; Wang *et al*. 2023; Yao *et al*. 2023).

Here, we developed an iterative growth model (IGMF) for WPC growth based on the IGM. Using observational data on global WPC carbon budgets (Fig. S1), we validated the IGMF and derived equations for WPC carbon budgets. These results improve our understanding of WPC carbon budgets and allow us to infer the maximum biomass (*M*_max_) of global WPCs based on their current carbon budget status. We also established causal links between global *M*_max_ and climate, which was used to project the direct effects of future climate change on *M*_max_.

## Box.1

The IGM is also built upon MST and further assumes that metabolism exponent (*b*) is not constant and the growth rate is constant during the time required to form a unit of tissue (*T*). Assuming that the total energy produced by respiration during *T* is used mainly used for the growth metabolism of new biomass (*f*(*m*)) and the maintenance metabolism of existing biomass (*m*), we can establish the following equation describing the growth rate (*f*(*m*)/*T*) can be derived (Shu *et al*. 2021):

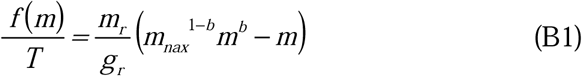

where *g_r_* and *m_r_* represent the stable respiration cost required to produce a unit of tissue and the variable maintenance respiration rate per unit of tissue, respectively; *m_max_* is the maximum biomass. Compared to the classical metabolic growth equation, the introduction of *T* reveals an iterative growth mechanism and two implicit constraints. The first is the total growth time (TGT) = *g_r_*/*m_r_* × (2*b* + 2)/(1 − *b*), which can be derived from the integral transform from *f*(*m*) to *m_max_.* The second is that *T* < *g_r_*/*m_r_*, which is determined by the thermodynamic significance of respiration. To counteract natural degradation (entropy increase), organisms must continuously use negative entropy to maintain the complexity, variety, and order of their components. During time *T*, the growth energy proportional to *g_r_* decreases the entropy of a new tissue unit relative to that of its free precursor monomers (Clarke 2019). Meanwhile, the maintenance energy, which is proportional to *Tm_r_*, maintains a low entropy state of an existing tissue unit and indicates its entropy accumulation during this time. Assuming that the old and new tissue units are identical, the synthesis of a new tissue unit is only possible if *Tm_r_* is less than *g_r_*; that is, *T* < *g_r_*/*m_r_*. As *T* approaches 0 and *g_r_*/*m_r_*, the integration or iteration of Eq. B1 yields two smooth functions driven by time (*t*), the Richards and Gompertz equations (Shu *et al*. 2021) (Supplementary Information Appendix A). Mathematically, the actual growth dynamics do not have an explicit analytic solution in most cases and lie somewhere between these two equations.

## Extension of the IGM to woody plant community NPP

The combination of the IGM and the density effect (West *et al*. 1999) highlighted the mediating effects of the ratio of the maintenance respiration coefficient to growth respiration coefficient, *m_r_*/*g_r_* (Box.1), WPC maximum biomass (*M*_max_), and current biomass (*M*) on WPC growth (see Supplementary Information, Appendix A for the derivation.).

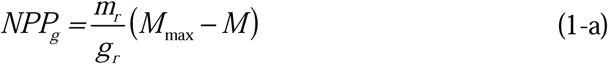

This framework was also true for structural net primary productivity (NPP_s_), which is mainly composed of biomass growth (NPP_g_) and leaf and fine root turnover (NPP_t_). By introducing a parameter *c* related to turnover, Eq. 1-a can be further expressed as.

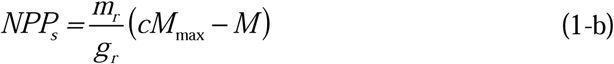

where WPC organ turnover rate (NPP_t_) is calculated as the difference between Eq. 1-b and Eq. 1-a, (*c* − 1) × *m_r_*/*g_r_* × *M*_max_ or NPP_s_/*M* + *m_r_*/*g_r_* × *M*/*c*. According to the growth– maintenance respiration paradigm, the autotrophic respiration rate during the WPC growth period (Ran) is *m_r_M* + *g_r_*NPP_s_, and the gross primary productivity required, GPP_s_, is equal to *m_r_M* + (1 + *g_r_*)NPP_s_.

Eq. 1-b also has two time-based expressions when *T* converges to its upper (*g_r_*/*m_r_*) (Box.1) and lower (0) bounds (see Supplementary Information, Appendix A for the derivation):

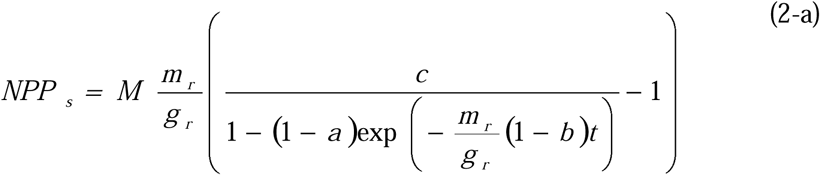

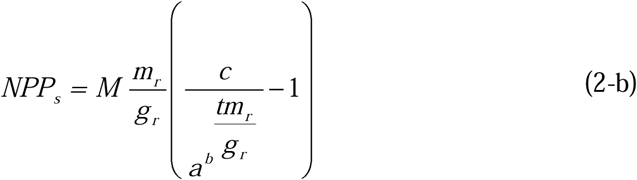

where *a* is the ratio of the initial WPC biomass *M*_0_ (where self-thinning started) to *M*_max_. We refer to Eqs. 1-b, 2-a, and 2-b together as the iterative growth model at the forest scale (IGMF). Mathematically, NPP_s_ lies between the thermodynamic lower and upper boundaries of the IGMF, as presented in Eqs. 2-a and 2-b. We labeled these as IGMF-L and IGMF-U, respectively. The IGMF also holds for aboveground woody productivity (NPP_aws_) driven by aboveground woody biomass (*M*_aw_), where the theoretical value of *c* is closer to 1. Theoretically, *m_r_* or *m_r_*/*g_r_* is temperature dependent and follows the Arrhenius equation (Thornley & Cannell 2000; Piao *et al*. 2010).

## IGMF fitting, evaluation and extension

We first fitted NPP_s_ and NPP_aws_ distributed across broad environmental gradients using Eqs. 1-a and 1-b, respectively. The results showed that the IGMF can explain most of the variation in these productivity, with an adjusted R² > 0.73 (*p* < 0.01) (Fig. 1A and **Fig. 2B**). Not only that, except for parameter *a*, both models yielded highly consistent fit parameters for the same dataset (Table. S1). This indicates that the error introduced by forcing the boundary to fit the data only affects parameter *a*, and that the other fitted parameters (repeatedly used in subsequent analyses) are accurate and robust. To further evaluate the IGMF and fitted parameters, we calculated NPP_s_ and NPP_aws_ for the WPCs in the different regions using these established equations, and compared the results with the observed values. On a logarithmic scale, the differences and ratios between predicted NPP_s_ (or NPP_aws_) and actual NPP_s_ (or NPP_aws_) were normally distributed with mean values of 0 and 1, respectively (Fig. S2), indicating an isometric relationship with a slope of 1 (Fig. 1C and Fig**. 1**D). These tests support the IGMF.

**Fig. 1.**
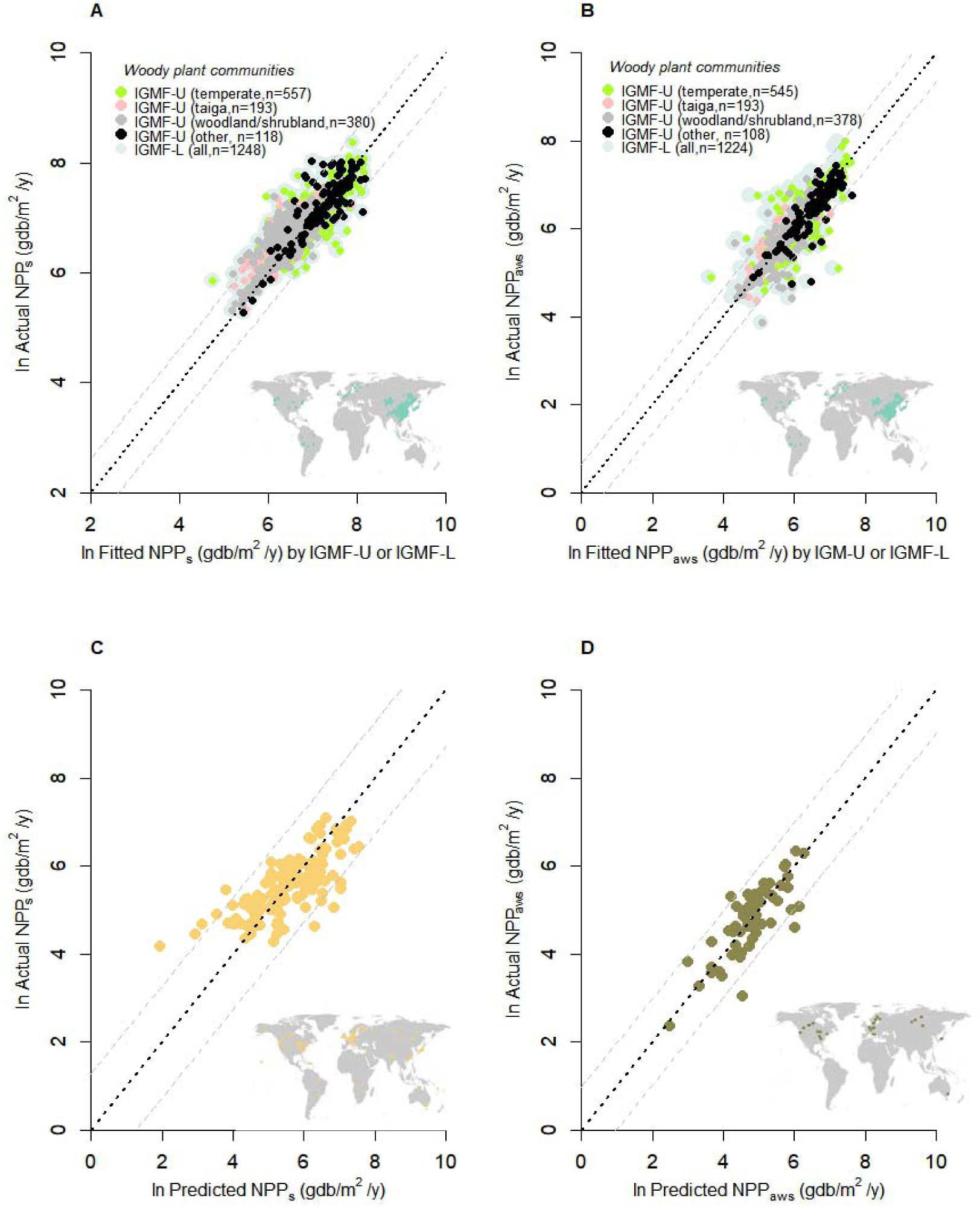
Fitting of IGMF-U (A) and IGMF-L (B) to structural WPC productivity, and prediction of NPP_s_ in other regions using these equations and the same fitting parameters (C and D) The black dashed line is *y* = *x*. The grey dashed line is 0.95 confidence interval. The colored scattered dots on the world map in the lower right corner of each figure are the WPCs involved in each analysis. C and D: The predictions are the average of the calculations by IGMF-U and IGMF-L

**Fig. 2.**
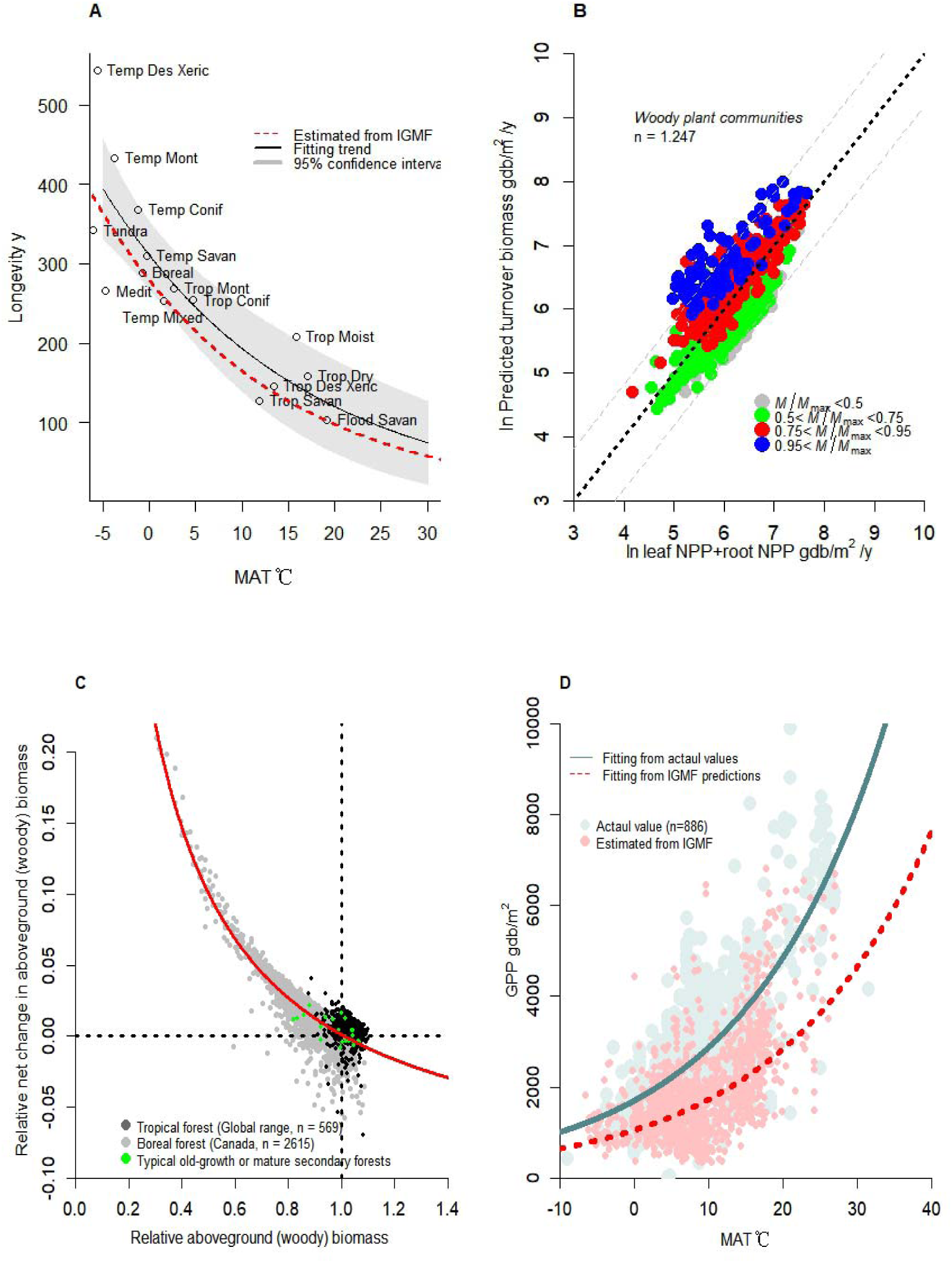
Comparison of predicted and actual results for (A) tree longevity, (B) WPC turnover, (C) forest biomass dynamics, and (D) gross primary productivity. A: Data are taken from ref. (Brienen *et al*. 2020). The red dashed line represents the average estimate based on the fitting parameters of Eq. 1-a (IGMF-U) and 1-b (IGMF-L), given by 14/exp (0.053x - 2.99), while in D, it is exp (0.05*x* + 6.96) (R^2^ = 0.30, *p* < 0.01). B: The black dashed line is *y* = *x.* The grey dashed line is 0.95 confidence interval. C: The red curve is log (1/0.105*x* − 0.90) (R^2^ = 0.81, *p* < 0.01). D: The green curve is exp (0.053*x* + 7.44) (R^2^ = 0.33, *p* < 0.01). The 95% confidence intervals for the exponential coefficients of the exponential functions in the figure are 0.047-0.057 (green solid line) and 0.045-0.054 (red dashed line).

We then validated the fitted *m_r_*/*g_r_*-MAT relationship. Given that the risk of tree mortality significantly increases with size or age (Brienen *et al*. 2020), we can assume TGT is approximately equal to tree lifespan. Extending *m_r_*/*g_r_* to TGT (14/exp(0.052MAT - 2.99)), we found that the estimated TGT-MAT curve is slightly lower than the temperature-lifespan relationship observed at the biome scale (Brienen *et al*. 2020) (Fig. 2A) and falls within its 95% confidence interval. This result supports the IGMF estimate of the *m_r_*/*g_r_*-MAT relationship.

Since NPP_t_ can be derived from Eq. 1-b containing the parameter *c*, we can assess the accuracy of *c* estimated in the above fit by comparing the predicted and observed values of NPP_t_. Leaf and root (organ) NPP_s_ consist of both their (its) growth (NPP_g_) and turnover (NPP_t_) components. As the growth allocation of the NPP_s_ decreases with stand age, the NPP_t_ becomes closer to the NPP_s_. This also means that when *M*/*M*_max_ is large, the predicted (organ) NPP_t_ and the observed organ NPP_s_ could maintain an isometric relationship with a slope of 1. Two data sets belonging to the classes 0.75 < *M*/*M*_max_ < 0.95 and *M*/*M*_max_ > 0.95 confirm this speculation. The fitted slopes (95% confidence intervals: 0.93 − 1.04 and 0.87 − 1.19) and intercepts (95% confidence intervals: −0.57 − 0.16 and −1.8 − 0.05) were not significantly different from 1 and 0, respectively (Fig. 2B). In addition, we constructed an indicator called the structural carbon potential (*ψ*NPP_s_) to measure the carbon sink capacity of WPCs, defined as NPP_s_/*M*. The minimum *ψ*NPP_s_ is equal to *m_r_*/*g_r_* (*c* − 1) according to Eq. 1-b, which is exactly the lower bound of the distribution of actual *ψ*NPP_s_ (Fig. S3). These tests verify the parameter *c* and highlight its ecological significance.

The IGM allows us to calculate the *M*_max_ of a WPC from its NPP_s_, *M*, and MAT. Since *M*_max_ represents a dynamic equilibrium between WPC growth and mortality, the relative net growth rate (RNG), defined as (growth - mortality)/biomass, is expected to be reach 0 when *M*/*M*_max_ equals 1. We analyzed the relationships between *M*/*M*_max_ and RNG in tropical forests, North American forests, and some old-growth or mature secondary forests. The results showed that RNGs gradually decreased as *M*/*M*_max_ increased, converging to 0 as *M* approached *M*_max_. This confirms the IGMF estimate of *M*_max_ (Fig. 2C).

We also calculated the GPP_s_ and found that at a growth efficiency of 0.75 or *g_r_* = 0.33 (Thornley & Cannell 2000), it has the same temperature sensitivity as actual GPP. This indicates a stable ratio of 0.58 (or 0.37) between nonstructural carbohydrates (NPP_n_) (i.e., GPP − GPP_s_) and GPP_s_ (or GPP), which allows NPP_n_ to be expressed as:

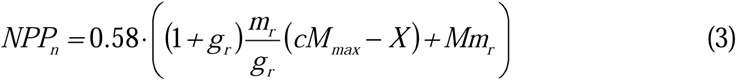

NSC allocation to WPC autotrophic respiration during nongrowing stages and other processes, such as allocation to fungi, root exudation, heartwood formation, secondary metabolite production, and embolism repair. These processes typically account for more than 25% of GPP (Smith & Smith 2011). In the long term, NPP_n_ (Eq. 3) may not accumulate continuously and should be roughly equal to the current size of the NSC pool. Using Eq. 3, we calculated the NSC pool size along the biomass gradient. The actual data (Furze *et al*. 2019) support this calculation (Fig. S4). These fits and tests (**Fig. 2**) strongly support the generalisability of the IGMF across broad environmental gradients, implying that other carbon budget processes also follow IGMF extensions (see Supplementary Information, Appendix B).

## Global WPC maximum biomass

Using the IGMF (Eq. 1-b) and raster data, we calculated the mean *M*_max_ for global WPCs at a resolution of 10 km × 10 km from 2016 to 2020, and validated these estimates with observations. First, the 95th percentile of current biomass for different forests within the same terrestrial biome represents their maximum biomass under ideal conditions, aligning with our estimated 95th percentiles of maximum biomass (Fig. 3A). Second, based on Eqs. S5 and S6, the GPP equals NPP/0.63 when M equals *M*_max_, a prediction confirmed by GPP and NPP data for WPCs with *M* close to *M*_max_ (*M*/*M*_max_ > 0.90) (Fig. 3B).

**Fig. 3.**
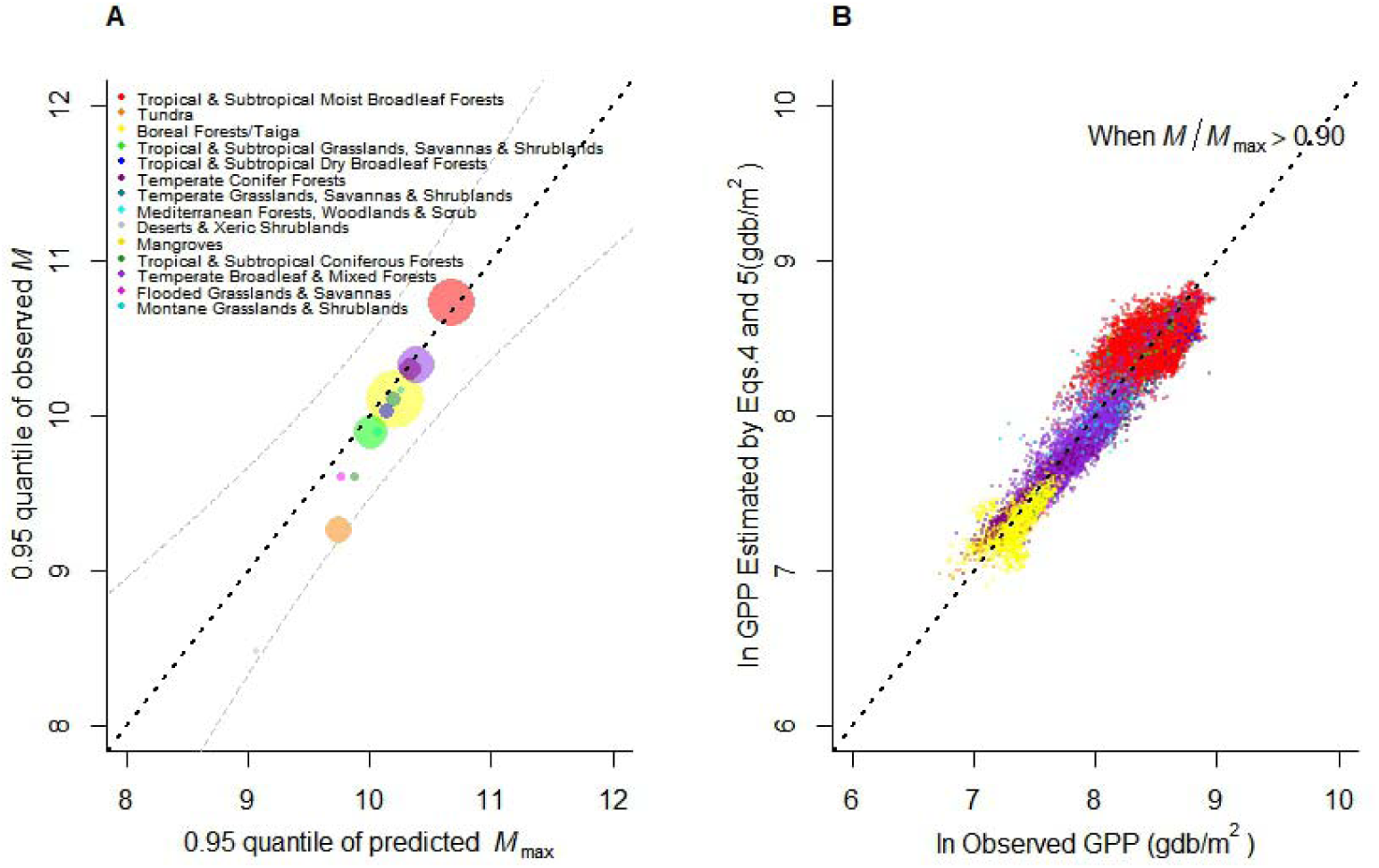
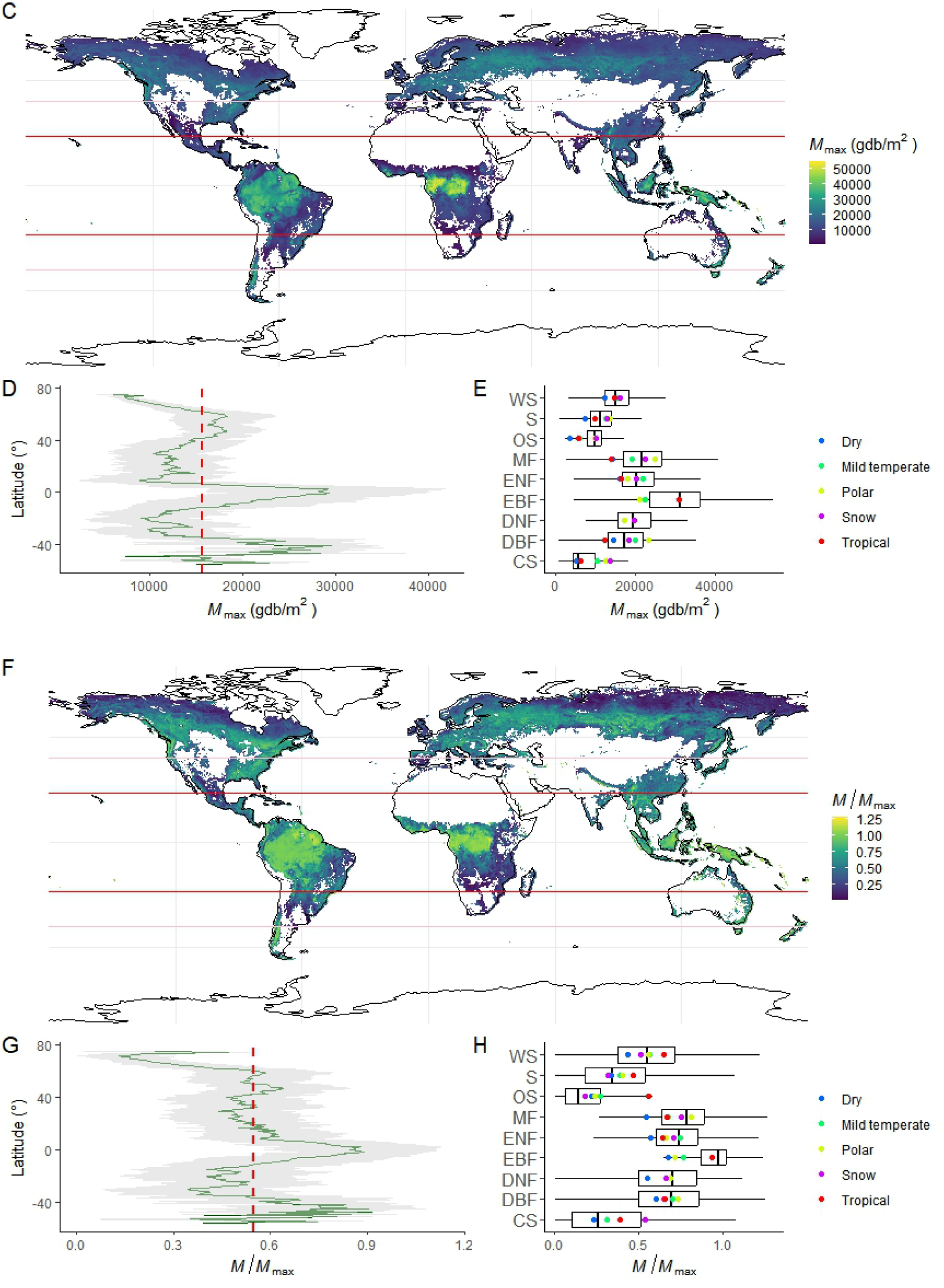
Estimated WPC maximum biomass vs. observed values (A), estimated WPC GPP vs. observed values (B), global WPC maximum biomass (C, D, E) and the ratio of current biomass to maximum biomass (F, G, H) A: The size of the points increases with the sample size. A and B: The black dotted line is *y* = *x* (R^2^ > 0.95, *p* < 0.01). The grey dashed line is 0.95 confidence interval. The meaning of the scatter colors in B is consistent with that in A. C and F: Solid red lines on the map indicate tropical boundaries and solid light red lines indicate subtropical boundaries. D and G: The red dashed line indicates the mean value. E and H: The abbreviations on the *y*-axis represent different WPCs, the meanings of which can be found in Fig. 4A. Coloured dots indicate the mean values.

Current WPCs have an *M*_max_ ranging from 6,349 to 35,111 gdb/m² (0.05 to 0.95 quartiles), with an average of 15,673 gdb/m² (Fig. 3C and Fig**. 3**D). Higher values (>35,111 gdb/m²) are mainly in low-latitude regions (Fig. 3C). Among forests, evergreen broadleaf forests exhibit the highest *M*_max_ (29,833 ± 9,629 gdb/m²), significantly exceeding other forest types, which range from 18,094 to 21,716 gdb/m² (Fig. 3E). The total *M*_max_ for evergreen broadleaf forests is also the highest among forests at 380 Pg, compared to mixed forests (174 Pg), evergreen needleleaf forests (81 Pg), deciduous needleleaf forests (6.7 Pg), and deciduous broadleaf forests (60 Pg). In shrublands, woody savannas have the highest *M*_max_ (15,803 ± 4,796 gdb/m²), while other shrublands range from 7,762 to 11,692 gdb/m² (Fig. 3E), which are much lower than forests. However, the total *M*_max_ for shrublands (750 Pg) surpasses that of forests (702 Pg), with woody savannas (271 Pg), tropical savannas (263 Pg), and open shrublands (212 Pg) being the largest contributors. *M*_max_ is significantly lower in dry regions than in other climate zones. By climate classification, *M*_max_ is significantly lower in dry regions than in others (Fig. 3E). Total *M*_max_ is highest in snow or continental/microthermal climates (678 Pg), compared to tropical (452 Pg), temperate (220 Pg), dry (73 Pg), and polar climates (29 Pg).

The *M/M*_max_ ratio, reflecting the realized growth potential of WPCs, shows a global pattern similar to that of *M*_max_, with significantly higher values near the equator (Fig. 3C and Fig**. 3**F). Globally, this ratio averages 0.64, with forests displaying a higher ratio (0.79 ± 0.10) compared to shrublands (0.41 ± 0.22) (Fig. 3H). Evergreen broadleaf forests have the highest ratio (0.96), while open shrublands have the lowest (0.21) (Fig. 3H). Based on this, we estimated the growth potential of WPCs. With a current biomass of 791 Pg, WPCs can grow by an additional 518 Pg, with forests and shrublands contributing 88 Pg and 430 Pg, respectively. Globally, _m_ost biomass growth will occur in the Northern Hemisphere, particularly above the subtropics (Fig. 3F and Fig**. 3**G). In snow or continental climates, WPCs could accumulate 294 Pg of biomass, more than in tropical (56 Pg) or temperate regions (73 Pg).

## Projected significant reduction in maximum biomass of evergreen broadleaf forests

Using extreme gradient boosting (XGBoost) models, we analyzed the effects of bioclimatic variables, the stable biomass-structure relationship (SBSR—defined as the residual between observed and predicted biomass-to-leaf area ratios), atmospheric CO_2_, soil properties (texture, saturated water content, organic matter content), and conservation strategies on *M*_max_. These factors effectively explained the variation in *M*_max_ among different WPC types (RMSE: 0.14 to 0.21, R^2^: 0.71 to 0.87) (Fig. 4A), with SBSR and soil saturated water content (wcsat), related to soil structural quality, significantly affecting *M*_max_ across all WPC types (Fig. 4B).

**Fig. 4.**
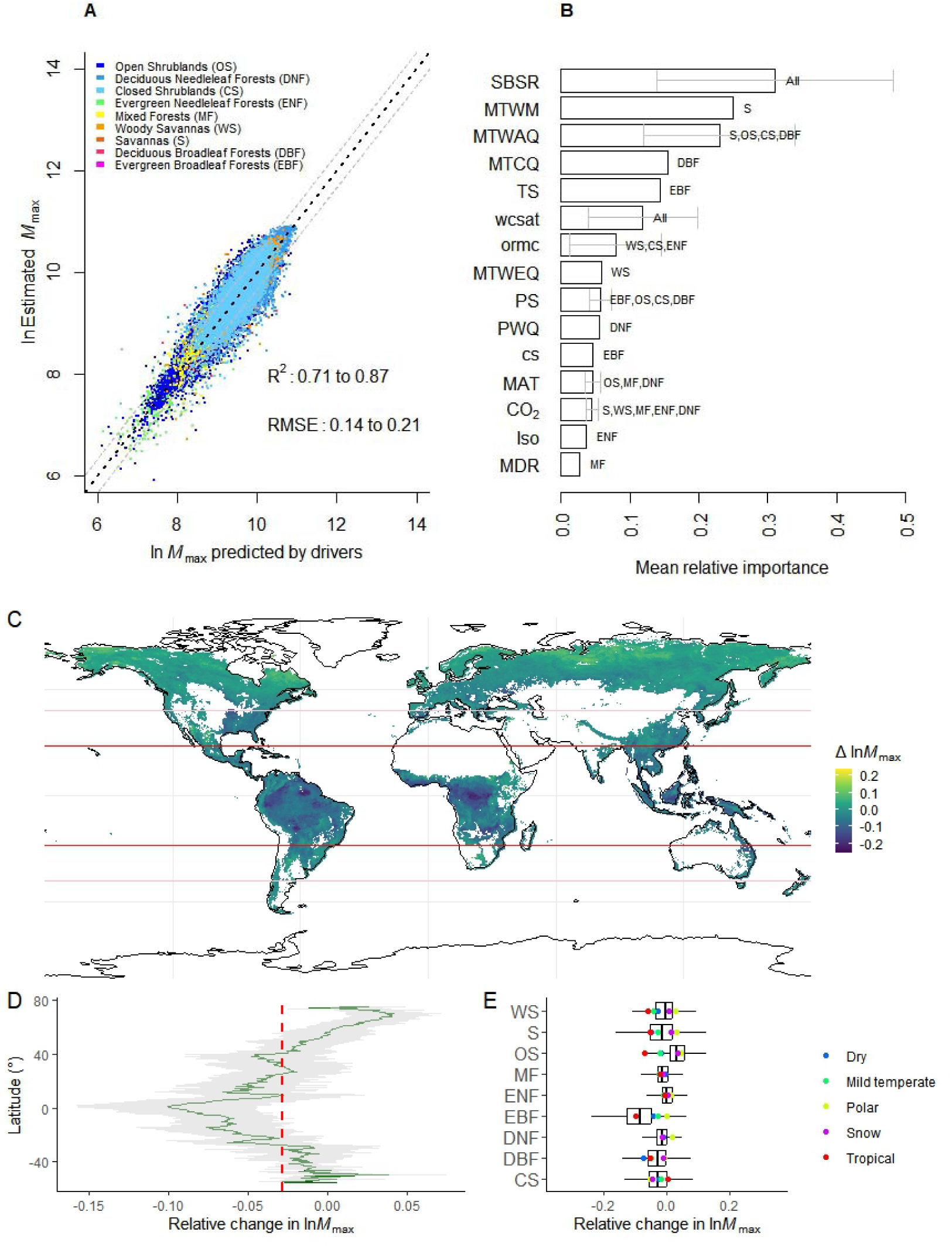
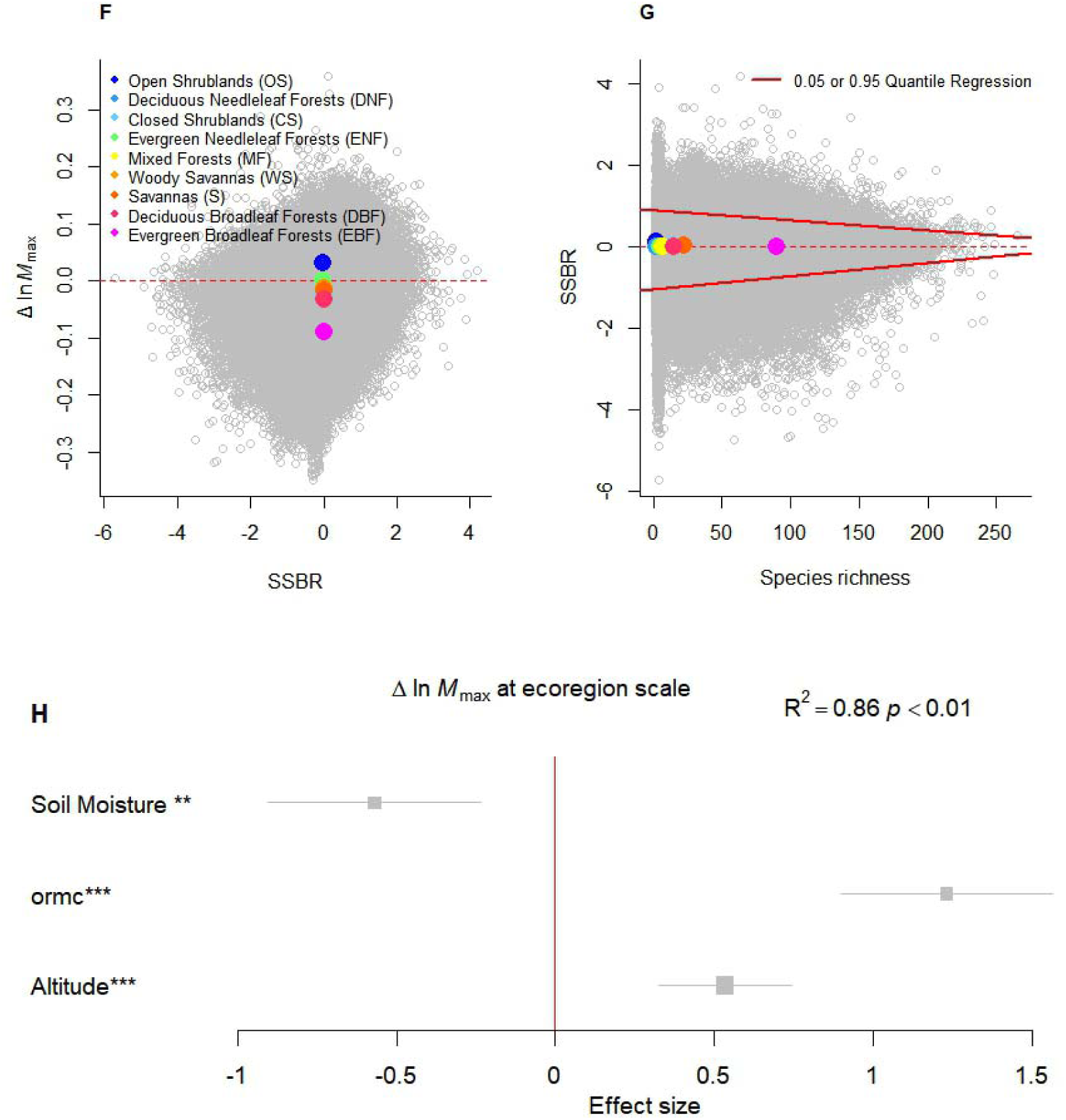
Machine learning assessment (A), relative importance of drivers (B), future relative changes in WPC maximum biomass (C, D, E), stability of this relative change (F, G) and its key drivers at the ecoregion level (H). A: The black dotted line is *y* = *x* (R^2^ > 0.71, *p* < 0.01). The grey dashed line is 0.95 confidence interval. B and H: SBSR, stable biomass-structure relationship; MTWM, max temperature of warmest month; MTWAQ, mean temperature of warmest quarter; MTCQ, mean temperature of coldest quarter; TS, Temperature Seasonality; wcsat, soil saturated water content; ormc, organic matter content; MTWEQ, mean temperature of wettest quarter; PS, precipitation seasonality (coefficient of variation); PWQ, precipitation of wettest quarter; cs, conservation strategies; MAT, mean annual temperature; Iso, Isothermality; MDR, mean diurnal range (mean of monthly (max temp - min temp)). The horizontal grey line is the standard deviation. C: Solid red lines on the map indicate tropical boundaries and solid light red lines indicate subtropical boundaries. Δln *M*_max_, relative change, defined as ln *M*_max (2100)_/ln *M*_max_ - 1, where *M*_max (2100)_ is the mean *M*_max_ from 2080 to 2100. D: The red dashed line indicates the mean value. E: The abbreviations on the *y*-axis rrepresent different WPCs, the meanings of which can be found in Fig. 4A. The colored dots in the lower right panel in C indicate the mean values. F and G: Colored dots represent the mean values of each WPC type within the coordinate system. The colors of the dots in G match those in F.

To capture the direct impact of climate on *M*_max_, we kept variables like plant leaf form, habit, and soil characteristics constant and used XGBoost models to predict global *M*_max_ under future climate conditions. The results indicate a decrease in total *M*_max_ with global warming (**Fig. S6**). By the end of the century, about two-fifths of WPCs will see an increase in *M*_max_, while three-fifths will see a decrease. The total *M*_max_ is projected to decrease by 266 Pg, mainly in the tropics (Fig. 4C and Fig**. 4**D). Evergreen broadleaf forests will have the largest reduction, decreasing by 60%, up to 228 Pg (Fig. 4E and Fig**. S7**). Among shrublands, savannas will decrease the most, with a total *M*_max_ reduction of 15% or 39 Pg. These decreases are mainly in tropical and subtropical regions like the Brazilian Plateau and the Mississippi Plain (Fig. 4C). In contrast, open shrublands will see a 34% increase in total *M*_max_, or 72 Pg, under both low and high climate warming scenarios (Fig. 4G and Fig**. S8**).

Further analysis reveals that future changes in *M*_max_ are related to species richness. In our regression models, *M*_max_ is influenced by both environmental factors and SBSR, which captures the remaining variation in *M*/*SLA* unexplained by stand age and environmental factors. As SBSR approaches 0, *M*_max_ becomes more likely to change (Fig. 4F). Interestingly, SBSR may be constrained by species richness (Fig. 4G), indicating that the greater the species richness is, the closer the SBSR is to 0 and the more sensitive *M*_max_ is, and vice versa, the more stable it is. However, at the ecoregional scale, relative changes in *M*_max_, whether increasing or decreasing, are independent of SBSR and are instead driven by soil moisture, soil organic matter content, and altitude (Fig. 4H and Fig**. S9**).

## Discussion

Plant growth, particularly tree growth, often shows complex, inconsistent, and variable trajectories influenced by genetics, climate change (Sheil *et al*. 2017; Begović *et al*. 2023), and uneven stand structure (Forrester 2019). However, our study supports the idea that, at the aggregate level, WPC growth follows a defined metabolic growth mechanism, which can enhance our understanding of WPC carbon sequestration. Some key findings include but are not limited to the following:

First, the carbon budgets of WPCs can be described by *λm_r_ M*(*γM*_max_/*M* - 1), where *λ* and *γ* are determined by *g_r_*, *c*, and specific carbon processes, and *M*_max_/*M* depends mainly on mr and stand age (Eqs. 2a and 2b). Physically, this reflects energy conservation: the autotrophic respiration rate of a forest (Ran = *m_r_M + g_r_*NPP_s_) equals its maintenance respiration rate at maximum biomass (including turnover biomass, *m_r_cM*_max_). Ecologically, this relationship shows that species composition, abundance, plant physiology, and inter- or intraspecific relationships influence growth by affecting *M*, *M*_max_, and *g_r_*/*m_r_*. For example, disturbances can decrease *M*, and warming can increase *m_r_*, both affecting the growth-mortality equilibrium (*M*_max_). These equations also show global convergence (Fig. 2Fig. 3), which may be attributed to the pipeline structure connecting leaves, stems, and roots, as well as the similarity of cellular components in plants. The pipeline structure ensures a linear allocation of NPP_s_ (Zhao *et al*. 2005; Malhi *et al*. 2011; Chen *et al*. 2019) between the crown, wood, and fine roots, related to tree self-thinning and parameter *c*, and similar cellular composition maintains stable *g_r_* and MAT-*m_r_* relationships. Thus, long-term changes in WPC growth or carbon budgets should follow the IGMF. For example, although CO_2_ fertilization can raise GPP rather than biomass in mature forests since most of the additional carbon is rapidly respired (Jiang *et al*. 2020), this decoupling or increase in GPP is still limited by the forest canopy or biomass. Forest biomass or stand age is still the main variable causing a decrease in carbon use efficiency (CUE, NPP/GPP) (DeLucia *et al*. 2007). Eventually, the CUE converges to 0.28, as predicted by the IGMF (Fig. S10).

Second, unlike projections from combined models (e.g., geobiochemical, spatial, and machine learning) (Zhu *et al*. 2018; Walker *et al*. 2022; Mo *et al*. 2023; Roebroek *et al*. 2023; Yu *et al*. 2024), the IGMF estimates of *M*_max_ do not require the integration of other factors and ecological processes but also offer accurate estimates. For example, our calculations estimate the near-term global total *M*_max_ (carbon stock) at 1,451 Pg db or 726 Pg C, which is close to that of Walker *et al*. (2022) at 796 Pg C and that of Mo *et al*. (2023) at 760 Pg C. The additional biomass growth or carbon storage potential in current WPCs is 518 Pg db (259 Pg C), with 17% in forests and 83% in shrublands, which aligns with the findings of Mo *et al*. (2023) at 226 Pg C and Walker *et al*. (2022) at 224 Pg C. In addition, the calculation results based on IGMF can visually reflect the probability of decline in WPCs. Some WPCs have *M*_max_ values lower than the current *M* values (Fig. 3F), indicating a risk of decline due to insufficient energy intake used for maintenance metabolism. The smaller the *M*/*M*_max_ ratio is than 1, the greater the likelihood of WPC decline. In contrast to some model-based expectations (Huntingford *et al*. 2013), we found that this decline is more likely in the tropics. Observations from the Amazon support this trend, showing that productivity gains have leveled off and that tree mortality has increased (Brienen *et al*. 2015). In fact, *M*_max_ represents the biomass at which growth and mortality offset each other under current conditions (**Fig. 2C**), varying with species, structure, environment, and management. When *M*/*M*_max_ > 1, the community cover is greater (0.96 _±_ 0.05) than when *M*/*M*_max_ < 1 (0.87 _±_ 0.10) (*p* < 0.01). In the former, plants compete more for resources and are more sensitive to disturbances (Brienen *et al*. 2015; Shu *et al*. 2019), making *M*_max_ vulnerable to decline. While the IGMF does not directly quantify these influences, its estimates reflect their effects on *M*_max_.

Third, in the long run, shrublands in the Northern Hemisphere still have high carbon sequestration potential. The projected maximum biomass for forests in China, Russia, the USA, and Canada from 2080--2100 was 12.4, 41.3, 28.6, and 28.4 Pg db, respectively, and for shrublands, it was 32.5, 397, 78.7, and 140 Pg db, respectively. These estimates are also accurate. For example, China’s WPCs are estimated at 22 Pg C, similar to that reported by Zhen *et al*. (2024) at 21.6--24.3 Pg C. The ratio of current biomass to future maximum biomass (2060--2080) in Canadian forests is 0.73, comparable to that reported by Zhu *et al*. (2019) at 0.78 ± 0.44. Collectively, these countries could accumulate 328.5 Pg of biomass between 2020 and 2100, covering two-thirds of the total accumulation gap. Additionally, evidence suggests that global conditions are shifting toward those historically favoring deciduous forests over evergreen forests (Ma *et al*. 2023), potentially further diminishing the carbon sequestration benefits of forests.

Fourth, species richness reduces species stability of biomass per unit leaf area (i.e., stable *M*/*SLA*, SSBR), enhancing *M*_max_ responsiveness to climate change, where the response direction of *M*_max_ to future climate change depend on soil moisture, organic matter, and altitude. In our regression models, SSBR is an indicator that depends only on species and their composition, dominating the stability of *M*_max_. Since high species richness typically reduces species synchrony (Valencia *et al*. 2020), the composition of species in such communities is more likely to shift with changing climate conditions, weakening the relationship between SSBR and species, and thereby reducing the stability of *M*_max_. However, species richness alone does not determine *M*_max_’s specific response to climate change. Under global warming, the current soil organic matter content primarily influences the direction of this response (Fig. S9). Higher soil organic matter content increases the probability of *M*_max_ rising; lower content increases the probability of it falling. Consequently, evergreen broadleaf forests, which have the highest species richness and the lowest soil organic matter content, show the greatest decline in *M*_max_ (Table. S3). Although the future global pattern of *M*_max_ is also shaped by bioclimatic variables, a series of belowground physiological and ecological processes related to organic matter accumulation (e.g., increased root exudates, arbuscular mycorrhizal formation and activity, and soil respiration) will become especially important.

In summary, our study provides a theoretical framework for understanding the growth of woody plant communities (WPCs) and their carbon sequestration potential. Based on a global analysis, we reveal the role of species richness and soil organic matter content in shaping WPC responses to climate change. These findings improve predictions of future carbon sequestration under changing global conditions.

## Glossary of main symbols

| Symbol | Meaning | Unit<sup>*</sup> |
| --- | --- | --- |
| $GPP$ | Gross primary productivity | $\text{g m}^{-2}\text{y}^{-1}$ |
| $NPP$ | Net primary productivity | $\text{g m}^{-2}\text{y}^{-1}$ |
| $NPP_n$ | Nonstructural carbohydrate | $\text{g m}^{-2}\text{y}^{-1}$ |
| $NPP_g$ | Biomass growth rate | $\text{g m}^{-2}\text{y}^{-1}$ |
| $NPP_t$ | Turnover rate (leaf and root) | $\text{g m}^{-2}\text{y}^{-1}$ |
| $NPP_s$ | $NPP_t + NPP_g$ | $\text{g m}^{-2}\text{y}^{-1}$ |
| $NPP_{as}$ | Aboveground $NPP_s$ | $\text{g m}^{-2}\text{y}^{-1}$ |
| $NPP_{aws}$ | Aboveground woody growth rate | $\text{g m}^{-2}\text{y}^{-1}$ |
| $M$ | Woody plant biomass | $\text{g m}^{-2}\text{y}^{-1}$ |
| $M_a$ | Aboveground biomass | 1 |
| $M_{aw}$ | Aboveground woody biomass | y |
| $M_{max}$ | Potential maximum $M_a$ | kg |
| $M_{amax}$ | Potential maximum $M_{aw}$ | kg |
| $M_{awmax}$ | Potential maximum $M_{aw}$ | kg/y |
| $M_0$ | Minimum biomass of woody plant communities where self-thinning occurs | kg/y |
| $a$ | $M_0/M_{max}$ | 1 |
| $b$ | Metabolic exponent, taken here as 0.75 | 1 |
| $c$ | Turnover-related parameter | 1 |
| $T$ | The formation time of unit tissue is primarily controlled genetically and by physiological activities, with the intrinsic or developmental growth rate independent of organism size | y |
| $m_r$ | Rate of maintenance respiration per unit of tissue | $\text{mg g}^{-1}\text{y}^{-1**}$ |
| $g_r$ | Cost of respiration needed to produce a unit of tissue | $\text{mg g}^{-1**}$ |
| $f(m)$ | Organism/plant mean growth increment during time $T$ at a given $m$ | g |
| $f_a$ | mean net growth rate of woody plant communities | g/y |
| $m$ | Organism/tree current size | g |
| $m_0$ | Organism/tree initial biomass | g |
| $o$ | Threshold biomass for growth | g |
| $m_{max}$ | Organism/tree potential maximum size/biomass | g |
| $Ra$ | Autotrophic respiration rate of woody plant communities | mg/y** |
| $Ran$ | Autotrophic respiration for growth and organ turnover in woody communities | mg/y** |
| $RNG$ | Relative net growth rate of woody plant communities, i.e., (growth – mortality)/biomass | 1 |
| $RNGa$ | Relative net rate of aboveground biomass growth, i.e., (aboveground growth – aboveground mortality)/aboveground biomass | 1 |
| $RNGaw$ | Relative net rate of aboveground woody growth, i.e., (aboveground woody growth – aboveground woody mortality)/aboveground woody biomass | 1 |
| TGT | Total growth time | y |

\* Units of time, respiration, and biomass are expressed as y (year), mg (carbon dioxide), and g (dry weight), respectively. \*\* mg carbon dioxide was converted to grams of biomass assuming biomass is 50% carbon.

## Acknowledgments

We thank Mr. Zhiqiang Xiao and Professor Lei Chen of Sichuan University for their valuable suggestions on the manuscript, and all researchers who provided accessible data.

## Author contributions

Conceptualization: SS

Methodology: SS

Investigation: SS, TX, YZ

Visualization: SS

Funding acquisition: XW

Project administration: XW, WZ

Supervision: XW, WZ

Writing – original draft: SS, KG

Writing – review & editing: TX, KG, XW, WZ, WW, XZ, ZH.

## Data and materials availability

Forest data are available at https://doi.org/10.1038/nature13470,

https://doi.org/10.5061/dryad.8bg44b0,

https://onlinelibrary.wiley.com/doi/10.1002/ecy.2229/suppinfo, and https://dx.doi.org/10.5521/forestplots.net/2020_2

Code will be made available on request.

## Supplementary Materials

### Materials and Methods

#### IGMF fitting and validation

The forest data we used to fit and test the IGMF (Eqs. 1-a and 1-b) covered most forest areas worldwide and comprised five primary datasets, as shown in Fig. S1. The first dataset we used was compiled by ref. (Michaletz *et al*. 2014). The authors summarized the growth and biomass of woody material, leaves, roots, tree age, and corresponding climate factors for 1,247 woody plant communities across broad climate gradients (green dots in Fig. S1). The second and third datasets were from ref. (Chen *et al*. 2016) or (Luo *et al*. 2019) and ref.(Sullivan *et al*. 2020). Both datasets recorded aboveground (woody) biomass, growth, and mortality dynamics of boreal (Canada, *n* = 2615, blue dots in Fig. S1) and tropical (global, *n* = 569, red dots in Fig. S1) forests during 1958-2019, as well as climate drivers. The fourth dataset was obtained from the open-access Global Forest Carbon database (ForC) (Anderson-Teixeira *et al*. 2018). This dataset contains previously published records of field-based measurements of ecosystem-level C stocks and annual fluxes, as well as disturbance history and methodological information. We compiled forest GPP and NPP, including growth and/or leaf and root turnover, mortality, stand age, and MAT, from this database. Forest productivity, mortality and biomass were converted to g of biomass, assuming that biomass is 50% carbon. In addition, we extracted global tree lifespans at the biome scale from the results of ref. (Brienen *et al*. 2020) and showed their variation along the MAT gradient.

We first fitted the NPP_s_ and NPP_aws_ of the first high-quality data set using Eqs. 1-a (IGMF-L) and 1-b (IGMF-U), and evaluated the goodness-of-fit. With these equations and the resulting fit parameters, it was possible to calculate forest NPP_s_ and NPP_aws_ for the fourth dataset based on *M*, stand age, and MAT. To maximize the use of the data, we used *M*_a_ multiplied by 1.34 as an approximation for *M* to compensate for some missing values in *M*. The factor 1.34 was calculated as the average ratio from the existing *M* to *M*_a_. We then obtained the ratios and differences between the simulated and actual values after applying a logarithmic transformation and assessed their distributions (Fig. S2). If these distributions approximate normal distributions with means of 1 and 0, respectively, the simulation can be considered theoretically consistent with reality. All calculations and analyses were performed in R version 4.0.2, using the "nls" function for fitting.

We tested the predictions of the IGMF for tree longevity, NPP_t_, and *M*_max_. According to the IGM, tree lifespan is approximately equal to 14*g_r_*/*m_r_*. Using MAT-*m_r_*/*g_r_* relationship generated by the above fits, we calculated tree lifespan across the MAT gradient. We then compared the actual changes in tree lifespan along MAT at the biome scale (Brienen *et al*. 2020) to the estimated temperature-lifespan curve (Fig. 2A).

According to Eq. 1-b, NPP_t_ and *M*_max_ are equal to NPP_s_/*M* + *m_r_*/*g_r_*·*M*/*c* and NPP_s_/*c*·*g_r_*/*m_r_* + *M*/*c*, respectively. Using these equations, we first calculated the corresponding NPP_t_ and *M*_max_ for the first dataset and then compared the calculated NPP_t_ with the measured results. Using the “lm” function in R version 4.0.2, we determined whether there was a linear isometry between the logarithm of the estimated NPP_t_ and that of the actual value (Fig. 2B). We also constructed an indicator termed the net carbon potential (*ψ*NPP_s_) (defined as NPP_s_/*X*). The minimum *ψ*NPP_s_ is equal to the ratio of NPP_s_ to *M*_max_ and, in theory, can be expressed as *m_r_*/*g_r_* (*c* − 1). To test the accuracy of *c* and *m_r_*, we compared the actual minimum *ψ*NPP_s_ and *m_r_*/*g_r_* (*c* − 1). The actual minimum was derived using the 5^th^ quantile regression of an exponential function on *ψ*NPP_s_ along the MAT gradient (Fig. S3) with the “quantreg” package in R version 4.0.2.

The second and third datasets were used to test the prediction of Eq. 1-a or Eq. 1-b for *M*_max_. Depending on whether the mortality data in the different datasets included leaf turnover, we constructed the relative net rate of aboveground woody growth, i.e., (NPP_aws_ − aboveground woody mortality)/*M*_awmax_, RNGaw) and the relative net rate of aboveground biomass growth (i.e., NPP_as_ − aboveground mortality)/*M*_a_, RNGa) to explore their changes with *M*_aw_/*M*_awmax_ and *X*_a_/*X*_amax_ (Fig. 3C). The change trend in RNGa or RNGaw along *M*_a_/*M*_amax_ or *M*_aw_/*M*_awmax_ was used as a proxy for the relationship between the relative net rate of forest biomass growth (RNG) and *M*/*M*_max_. The theoretical equation is valid if RNG → 0 and *M*/*M*_max_ → 1. We first calculated *M*_awmax_ for tropical and boreal forests using the generated parameters (Table. S1) and Eq. 2-a or Eq. 2-b. For the boreal forest dataset containing *M*_a_, NPP_as_ and corresponding mortality, we used *M*_a_ = 1.06 × *M*_aw_ and NPP_as_ = 0.54 × NPP_aws_ to obtain their *M*_aw_ and NPP_aws_, as well as *M*_amax_, where the inflation factors were the actual ratios of these variables, summarized from the boreal forests in the first dataset. On this basis, we predicted their RNGaw or RNGa. We also focused on the RNGaw and *M*_aw_/*M*_awmax_ of some typical old-growth or mature secondary forests (data from ref. (Anderson-Teixeira *et al*. 2018), Table. S2). These forests have been subjected to natural and/or historical human disturbances.

With *g_r_* and *m_r_* known, we calculated the GPP_s_ for the forest in the first dataset and then compared the differences between the GPP_s_ and the measured GPP from the fourth dataset along the MAT gradient. Compared with the actual GPP, the GPP_s_ were lower but had the same temperature sensitivity (Fig. 3D). Although they are from different sites, this gap is likely filled by temporarily stored nonstructural carbohydrates (NPP_n_) (Eq. 3). It is also likely that NPP_n_ is mainly used for autotrophic respiration in the nongrowing stage, implying that NPP_n_ may be generally equal to the size of the NSC pool. We tested this hypothesis using a set of published data on whole-tree total NSC storage versus tree biomass from ref. (Furze *et al*. 2019) (Fig. S4). If this finding is confirmed, then the sum (Rat) of Ran and NPP_n_ should be close to Ra. Using measured values from the fourth dataset and calculated values based on the first dataset, we compared their differences across the MAT gradient (Fig. S5).

#### IGMF-based global WPC maximum biomass and its future changes

The data used to drive the IGMF for calculating global forest *M*_max_ were primarily sourced from the Google Earth Engine (GEE) platform. We first constructed raster layers of various raster data at a 1 km resolution spanning the years 2017-2020. These raster maps included net primary productivity (from the MOD17A3HGF dataset), forest aboveground biomass (from the CCI BIOMASS dataset), land cover (from the MOD13Q1 dataset), atmospheric CO_2_ concentration (from the OCO2_GEOS_L3CO2_MONTH dataset, NASA), and key climatic variables (monthly maximum and minimum temperatures from ERA5 and precipitation from the CHIRPS dataset). Forest aboveground biomass was then expanded to total biomass by multiplying it by 1.32. Meanwhile, the key climate variables were further converted to 19 bioclimatic variables using the “BioVars” function in R version 4.0.2. Next, we distinguished forest rasters (vegetation with a cover greater than 0.2) using land use classification (from the MCD12Q1 dataset) and further considered their biome classification/conservation strategy (from the RESOLVE Ecoregions 2017 dataset) (Dinerstein *et al*. 2017), forest type (from the MOD13Q1 dataset), species richness (Liang *et al*. 2022), and soil properties (including texture, soil moisture, saturated water content, and organic matter content from the HiHydroSoilv2_0 dataset). Finally, we obtained raster maps of the mean values of the 19 bioclimatic variables at 20-year intervals from 2020 to 2100 from WorldClim, with a resolution of 30 arcseconds or about 1 km.

We used the IGMF to convert raster layers of *M*, MAT, and NPP into the map of forest maximum biomass (*M*_max_) at 1 km resolution, using the “raster” package in R version 4.0.2. However, these raster NPP maps may or may not include NPP_n_. To address this, we first used Eqs. 5 and 9 to obtain curves of NPP and NPP_s_ along the MAT gradient based on actual observations *M* and estimated *M*_max_. Then, we distinguish between the cases where the NPP raster data contains NPP_n_ or not when calculating *M*_max_. Where the raster NPP data are closer to the NPP (greater than the mean of the theoretical NPP and NPP_s_ curves), Eq. S6 is used to estimate *M*_max_, and for the remainder, Eq. S10 is used. To test the accuracy of this data classification, we introduced a key assumption that if the NPP raster data are close to the NPP_s_, then the GPP raster data and the NPP raster data must necessarily conform to Eqs. S5 and S6. This is because if the NPP raster data contains NPP_n_, its corresponding GPP raster data necessarily conforms to Eq. S5, implying that the ratio of the NPP raster data to the GPP raster data is between 0.64 and 0.83. Tests supported this speculation (Fig. S11).

To capture the environmental effects on *M*_max_ as comprehensively as possible while making them easy to quantify, we considered the ratio of forest biomass to specific leaf area (*M*/*SLA*). This ratio is influenced by a combination of species, stand structure, stand development stage (with stand cover as a proxy), and the environment. Theoretically, by separating the effects of stand development stage and environment on the ratio, we could extract information about species and stand structure that is independent of the environment, referred to as the stable biomass-structure relationship (SBSR). This information can be approximated by regression residuals. As the SBSR is environmentally stable, establishing the relationships among the SBSR, the environment, and *M*_max_ fully captures the environmental effect.

The extreme gradient boosting model (XGBoost), a modern decision tree-based regression technique, was used to conduct machine learning of *M*/*SLA* and *M*_max_. We chose XGBoost for its ability to handle nonlinear interactions, outliers, collinear predictors, large datasets, categorical and missing data; for its processing speed; and for its ability to minimize overfitting (Elith *et al*. 2008; Balzotti *et al*. 2020). We first determined the effects of environment and stand development stage on *M*/*SLA* by creating XGBoost models for different types of forests and used the fitted residuals to represent the stable biomass-structure relationship (SBSR). Then, we assessed the relative importance of the SBSR and other drivers, such as bioclimatic variables, atmospheric CO_2_, soil properties (texture, saturated water content, and organic matter content), and conservation strategies on *M*_max_ by creating new XGBoost models for different WPC types. After logarithmic transformation of *M*/*SLA* and *M*_max_, 80% of the data were used for training, and the remaining 20% were used for validation. We implemented XGBoost using the R version 4.0.2 package XGBoost. Model tuning parameters are explained in detail in (Chen & Guestrin 2016) and included eta, gamma, max depth, min child weight subsample. We used the root-mean-square error (RMSE) and coefficient of determination (R^2^) as metrics of performance. Finally, we further assumed that forest type, soil properties, and atmospheric CO_2_ would remain relatively stable and estimated *M*_max_ based on the established XGBoost model and future climate drivers. These stability assumptions were made to highlight the direct effect of climate on *M*_max_. On this basis, we calculated the relative change in ln *M*_max_ (denoted as Δln *M*_max_), defined as ln *M*_max (2100)_/ln *M*_max_ - 1, where *M*_max (2100)_ is the mean *M*_max_ from 2080 to 2100. Additionally, we explained the stability mechanism of *M*_max_ and then analyzed its relative change drivers at the ecoregional scale using multiple linear regression (MLR), with potential drivers including changes in bioclimatic variables, altitude, current soil properties (texture, saturated water content, and organic matter content), atmospheric CO_2_, stand structure (LAI and stand cover). To avoid multicollinearity, variables with an absolute correlation value greater than were initially excluded, and employed a variance inflation factor (VIF) threshold of 5 to eliminate variables that still exhibited strong correlations in the results.

## Appendix A

### Upper and lower boundaries of the growth trajectory

When *T* → 0 and *g_r_*/*m_r_,* integrating or iterating Eq. B1 will produce two smooth functions driven by time (*t*), i.e., the Richards and Gompertz equations.

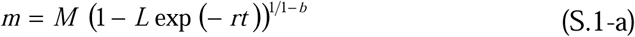

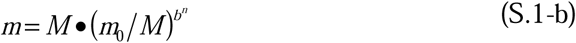

where *L* = 1-*M^b^*^-1^×*m*_0_, *r* = *m_r_*/*g_r_*(1-*b*), *m*_0_ is the first biomass observed, and *n* is the number of iterations and is equal to *t•m_r_/g_r_*. These results indicate that actual growth dynamics lie somewhere between these equations (Eqs. S.1-a and S.1-b) and may not be an explicit analytic solution in most cases.

### IGM for WPC structural NPP

Assuming that tree size heterogeneity is limited (e.g., normal distribution) and stand density (*d*) □ *m^-b^*(West *et al*. 1999), one can consider forest total growth as *f*(*m*)*d*. By simultaneously multiplying both sides of Eq. B1 by *d*, we can obtain:

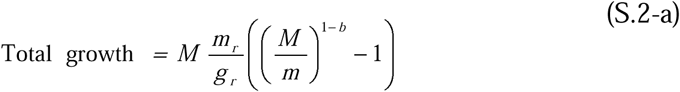

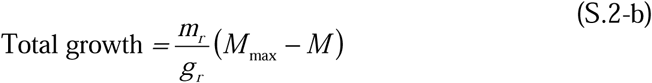

where *M*_max_ and *M* are the maximum biomass and current biomass of the forest, respectively. Forest carbon turnover is also an important component of NPP_s_ and can be incorporated into the same framework, resulting in

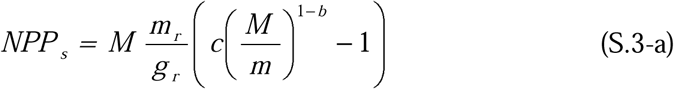

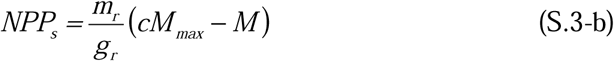

where *c* is a turnover-related parameter. Notably, the unit of biomass here is the same as that of NPP_s_ (structural NPP, including growth and organ turnover).

As *T* converges to 0 and *g_r_*/*m_r_*, the accumulation of *m* with time in the IGM (Eq. 1) conforms to the Richards and Gompertz Equations (Eqs. S.1-a and S.1-b), respectively (*9*). Because *M*/*M*_max_= (*m*/*m_max_*)*^1-b^*, we obtained the following:

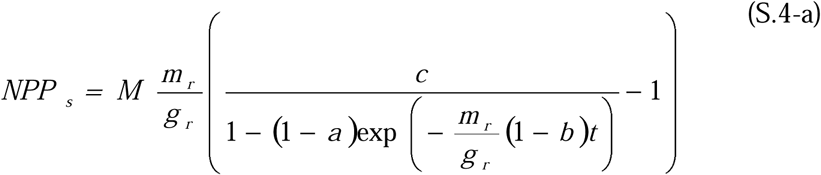

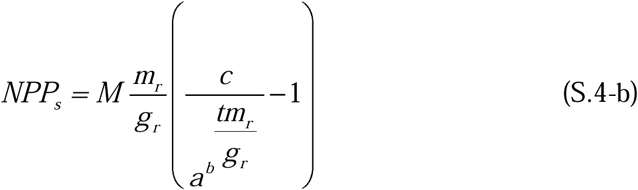

where *a* is the ratio of forest initial biomass to *M*_max_, *t* is forest age, and *b* is the metabolic exponent. Given that *g_r_*/*m_r_* is the upper limit of *T,* determined by the thermodynamic significance of respiration, we can term them the thermodynamic lower (IGMF-L) and upper (IGMF-U) boundaries of forest NPP_s_, and in theory, the real NPP is between the two boundaries.

## Appendix B

**Fig. 2** strongly support the generalisability of the IGMF across broad environmental gradients. Thus, we can reduce Eqs. 1-a, 1-b, 3, and their extensions to a set of kinetic equations for the global WPC carbon budgets:

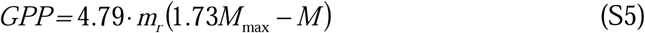

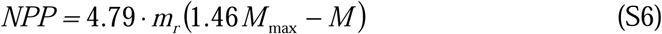

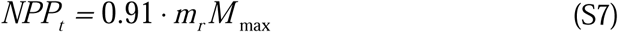

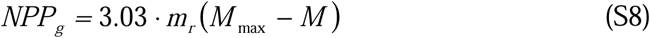

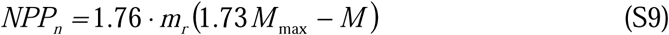

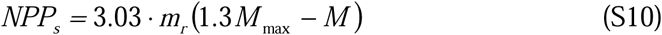

where changes in *M*/*M*_max_ over time (stand age) fall between the Richards and Gompertz equations (Eqs. S.1-a and S.1-b, respectively). The rate of autotrophic respiration necessary for WPC growth (Ran) is the difference between Eqs. S5 and S6, i.e., 1.3*m_r_M*_max_ and is proportional to the mean net growth rate (*f_a_* = 1/14 × *m_r_*/*g_r_* × *M*_max_) and the organ turnover rate (NPP_n_ = (*c* − 1) × *m_r_*/*g_r_* × *M*_max_). The total autotrophic respiration rate (Ra) is close to and does not exceed the sum of Ran and NPP_n_, i.e., 1.76*m_r_*(2.47*M*_max_ − *M*) (Fig. S5). These findings indicate that GPP, NPP_n_, and NPP_g_ decrease with WPC growth, but not NPP_t_. Overall, the total amounts of GPP, growth, autotrophic respiration during the growth period (excluding rhizosphere respiration), organ turnover, NPP_n_ and mortality (or coarse woody debris) were 25.1*M*_max_, 5.5*M*_max_, 6*M*_max_, 4.2*M*_max_, 9.1*M*_max_ and 4.5*M*_max_, respectively. Notably, these equations apply only after the WPC has undergone sustained self-thinning. Before this stage, variables such as GPP and NPP_g_ increase with biomass (Keeling & Phillips 2007).

## Appendix C

**Table S1.**
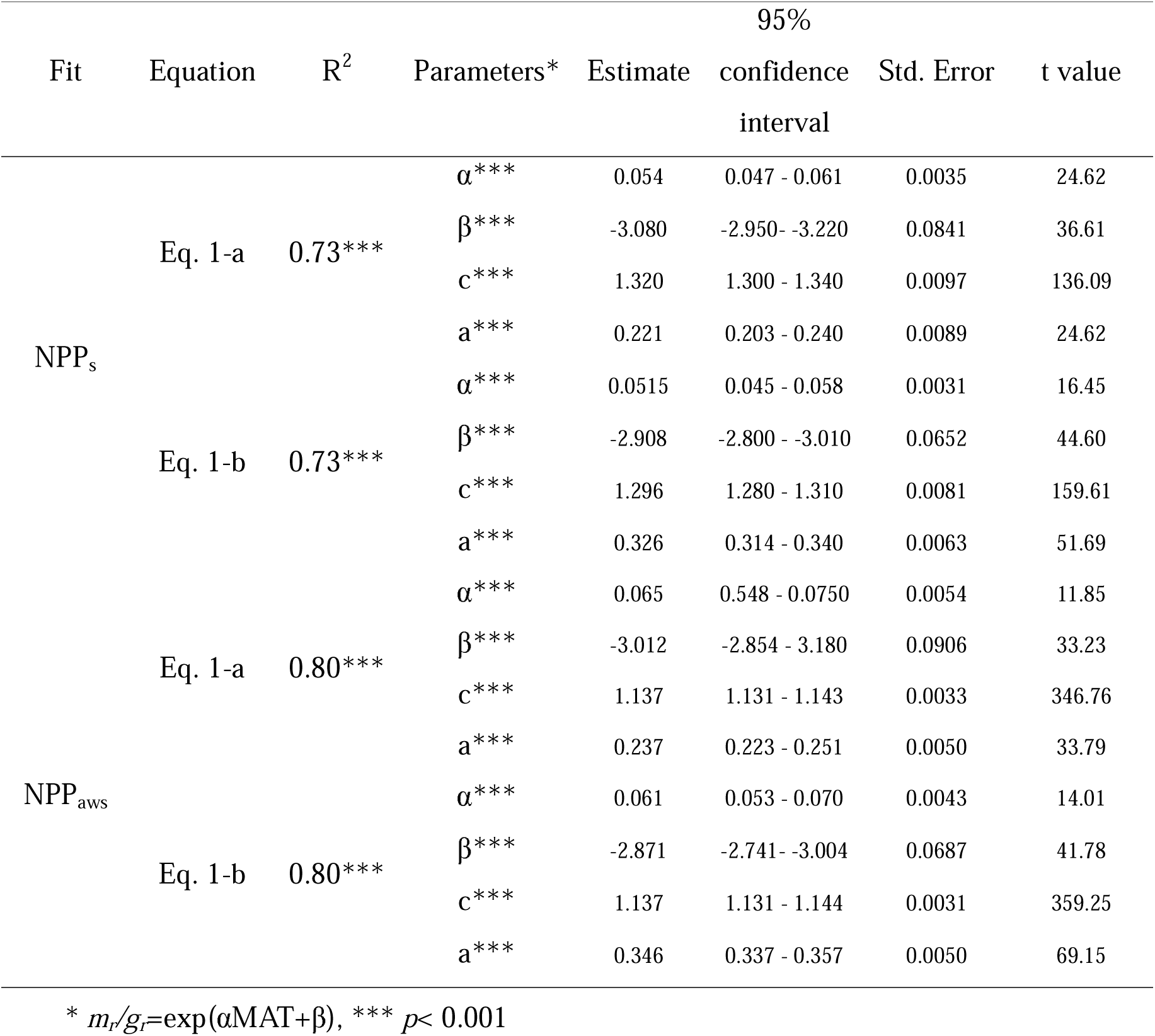
Eq. 1-a and/or 1-b fit parameters for NPP and aboveground woody plant growth.

**Table S2.** Machine learning results for *M*_max_: relative importance of top 5 drivers and model performance.

| WPC_Type | Variable | Importance | RMSE | R<sup>2</sup> | Optimal nrounds |
| --- | --- | --- | --- | --- | --- |
| Savannas | MTWM | 0.25 | 0.18 | 0.83 | 700+ |
|  | SBSR | 0.21 |  |  |  |
|  | MTWAQ | 0.17 |  |  |  |
|  | wcsat | 0.07 |  |  |  |
|  | CO<sub>2</sub> | 0.03 |  |  |  |
| Evergreen broadleaf forest | SBSR | 0.39 | 0.16 | 0.83 | 372 |
|  | TS | 0.14 |  |  |  |
|  | cs | 0.05 |  |  |  |
|  | wcsat | 0.04 |  |  |  |
|  | PS | 0.04 |  |  |  |
| Woody savannas | SBSR | 0.45 | 0.17 | 0.71 | 463 |
|  | wcsat | 0.06 |  |  |  |
|  | MTWEQ | 0.06 |  |  |  |
|  | CO<sub>2</sub> | 0.05 |  |  |  |
|  | ormc | 0.04 |  |  |  |
| Open Shrublands | MTWAQ | 0.39 | 0.15 | 0.87 | 487 |
|  | wcsat | 0.18 |  |  |  |
|  | SBSR | 0.06 |  |  |  |
|  | PDM | 0.05 |  |  |  |
|  | MAT | 0.05 |  |  |  |
| Closed Shrublands | wcsat | 0.29 | 0.21 | 0.86 | 121 |
|  | MTWAQ | 0.17 |  |  |  |
|  | ormc | 0.16 |  |  |  |
|  | SBSR | 0.06 |  |  |  |
|  | PS | 0.06 |  |  |  |
| Deciduous Broadleaf forest | SBSR | 0.25 | 0.16 | 0.84 | 173 |
|  | MTWAQ | 0.18 |  |  |  |
|  | MTCQ | 0.15 |  |  |  |
|  | PS | 0.08 |  |  |  |
|  | wcsat | 0.07 |  |  |  |
| Mixed Shrublands | SBSR | 0.51 | 0.14 | 0.82 | 291 |
|  | wcsat | 0.12 |  |  |  |
|  | MAT | 0.06 |  |  |  |
|  | CO<sub>2</sub> | 0.04 |  |  |  |
|  | MDR | 0.03 |  |  |  |
| Evergreen needleleaf forest | SBSR | 0.44 | 0.16 | 0.73 | 166 |
|  | wcsat | 0.08 |  |  |  |
|  | CO<sub>2</sub> | 0.06 |  |  |  |
|  | ormc | 0.04 |  |  |  |
|  | Iso | 0.04 |  |  |  |
| Deciduous needleleaf forest | SBSR | 0.43 | 0.14 | 0.73 | 42 |
|  | wcsat | 0.15 |  |  |  |
|  | PWQ | 0.06 |  |  |  |
|  | CO<sub>2</sub> | 0.05 |  |  |  |
|  | MAT | 0.03 |  |  |  |

See Fig.4 for variable meanings

**Table S3.** Soil organic matter content, relative and net changes in *M*_max_ for various WPC types.

| Type | ormc (g/kg) | $\Delta M_{\max}$ | Net change in total<br>$M_{\max}$ (PG) |
| --- | --- | --- | --- |
| Closed Shrublands | 12.54 | -0.22 | -0.84 |
| Deciduous Broadleaf Forests | 13.42 | -0.28 | -16.77 |
| Deciduous Needleleaf Forests | 22.49 | -0.14 | -0.934 |
| Evergreen Broadleaf Forests | 9.08 | -0.60 | -228 |
| Evergreen Needleleaf Forests | 20 | 0 | 0 |
| Mixed Forests | 18.54 | -0.13 | -22.62 |
| Open Shrublands | 28.6 | 0.34 | 72.08 |
| Savannas | 14.19 | -0.15 | -39.45 |
| Woody Savannas | 17.32 | -0.11 | -29.81 |

ormc: organic matter content
$\Delta M_{\max}$ : defined as $M_{\max(2100)}/M_{\max} - 1$ , where $M_{\max(2100)}$ is the mean $M_{\max}$ from 2080 to 2100.

**Fig. S1.**
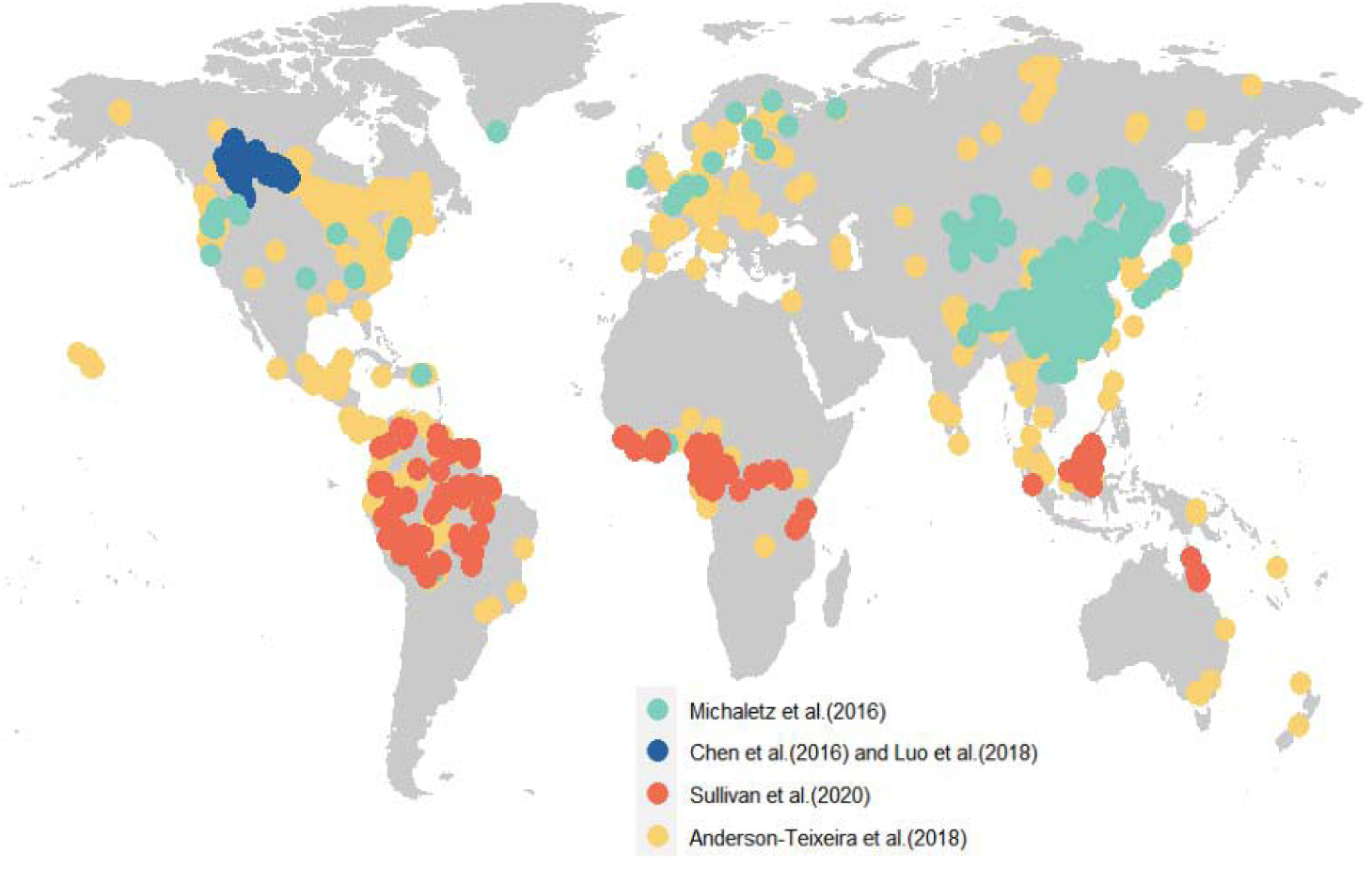
Geographical distribution of 7,140 compiled observations.

**Fig. S2.**
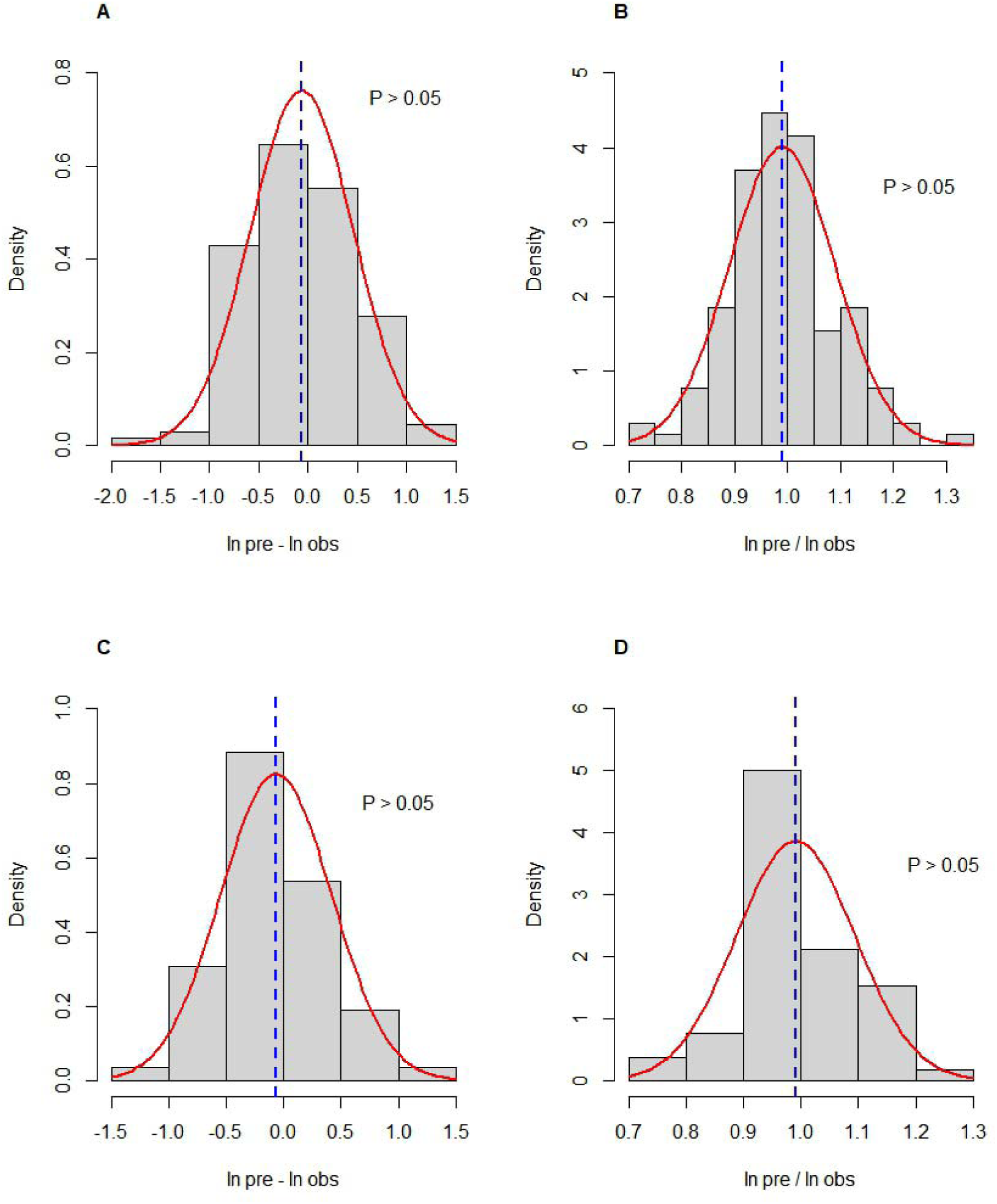
Distribution of the ratio and difference between the predicted and observed values for NPP_s_ (A and B) and NPP_aws_ (C and D) The red curve represents a normal distribution curve, and the blue dashed line represents the mean.

**Fig. S3.**
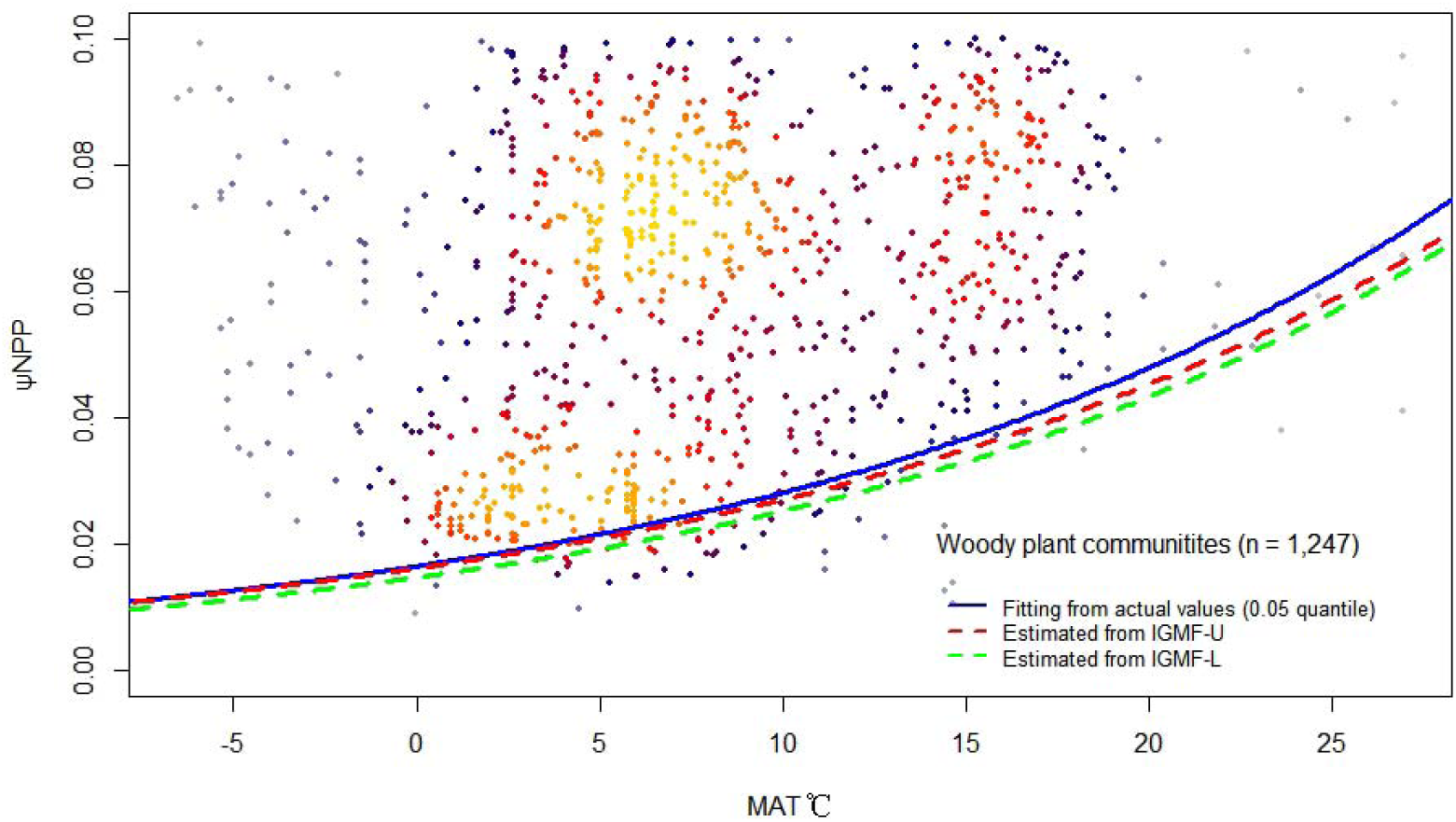
Changes in forest net carbon potential along the temperature gradient. The green line is exp(0.053MAT-4.10), derived from the 0.05 quantile fitting of mean annual temperature (MAT) to net carbon potential. Theoretical predictions according to the IGMF-L and the IGMF-U are 0.30exp(0.052MAT-2.91) (red dotted line) and 0.32exp(0.054MAT-3.08) (green dotted line), respectively.

**Fig. S4.**
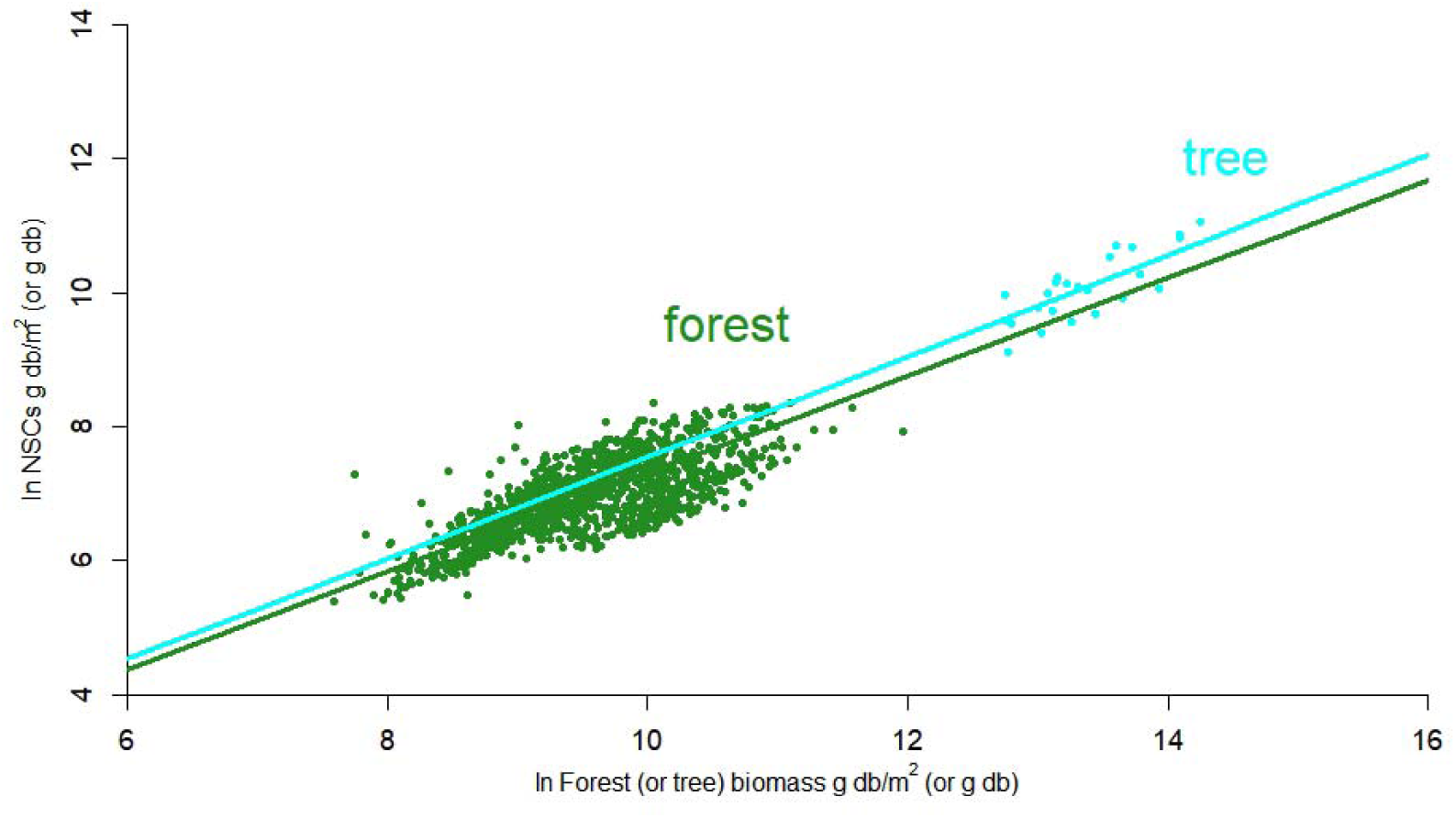
Model-predicted forest NSC pool size vs. forest biomass and actual tree NSC pool size vs. tree biomass. Using Eqs. 1-a and 1-b, we estimated the NSC pool size along the forest biomass gradient (data from ref. (Michaletz *et al*. 2014)), shown as green points. Actual changes in tree NSC pool size with tree biomass are shown as bright blue points (data from ref. (Furze *et al*. 2019), Fig. 2). A linear regression with no intercept was fitted to these data. Ordinary least squares (OLS) regressions of forest and tree data yielded *y* =0.73*x* and *y* = 0.75*x*, where the fitted slopes of the two equations were not significantly different (*p* > 0.05). Note that it was theoretically assumed that *y* = *x*, so no intercept was set.

**Fig. S5.**
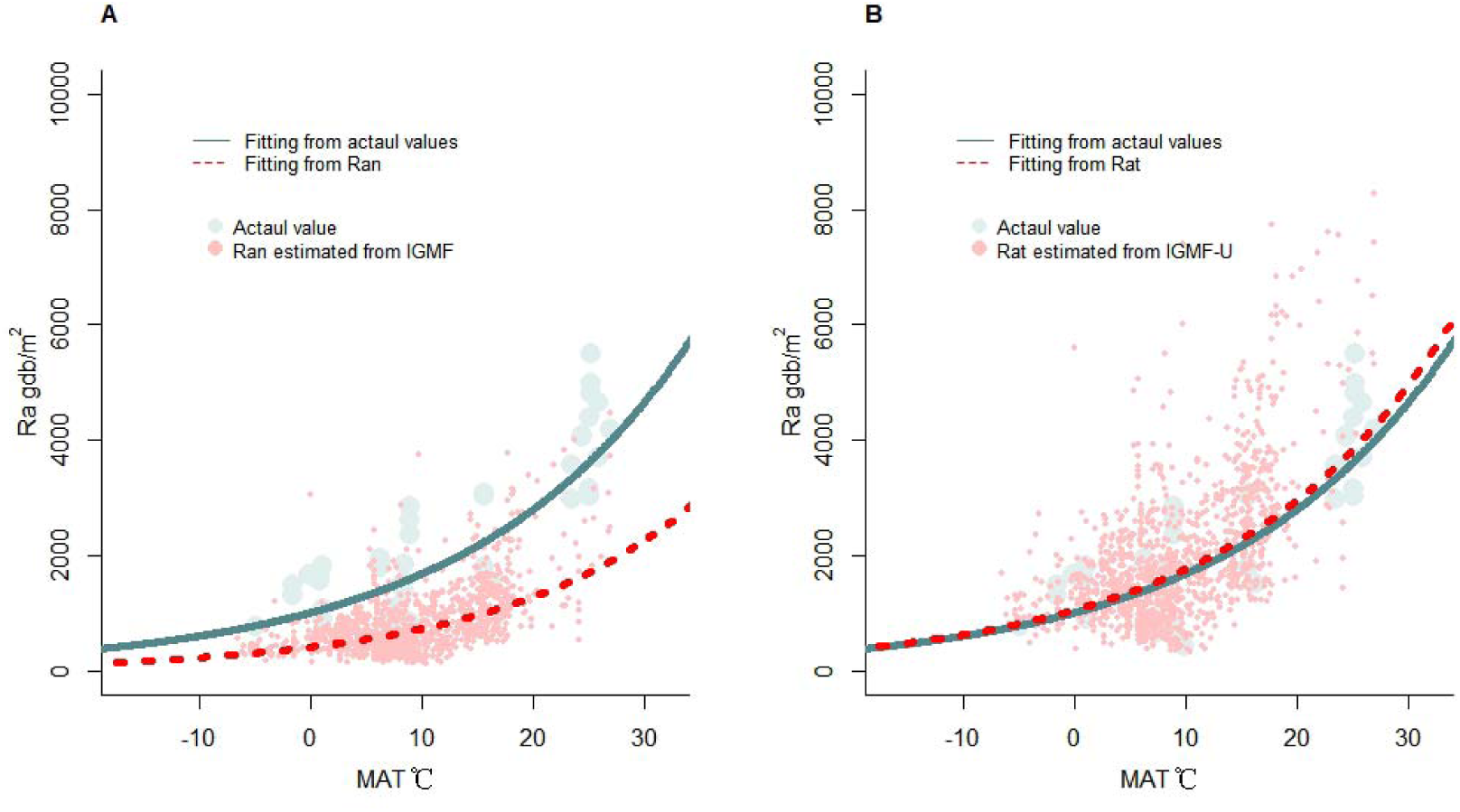
Comparison of actual respiration with respiration during growth (A) and comparison of actual respiration with the sum of respiration during growth and nonstructural carbohydrates (B) Ran: the sum of growth respiration and maintenance respiration during forest growth Rat: Ran + temporarily stored nonstructural carbohydrates (NPP_n_) The dark green line is exp(0.058*x* + 6.93) (R^2^ = 0.53, *P* < 0.01). The red dotted line in Fig. S5A is exp(0.056*x*+ 6.08) (R^2^ = 0.30, *P* < 0.01). The red dotted line in Fig. S5B is exp(0.050*x* + 7.03) (R^2^ = 0.30, *P* < 0.01).

**Fig. S6.**
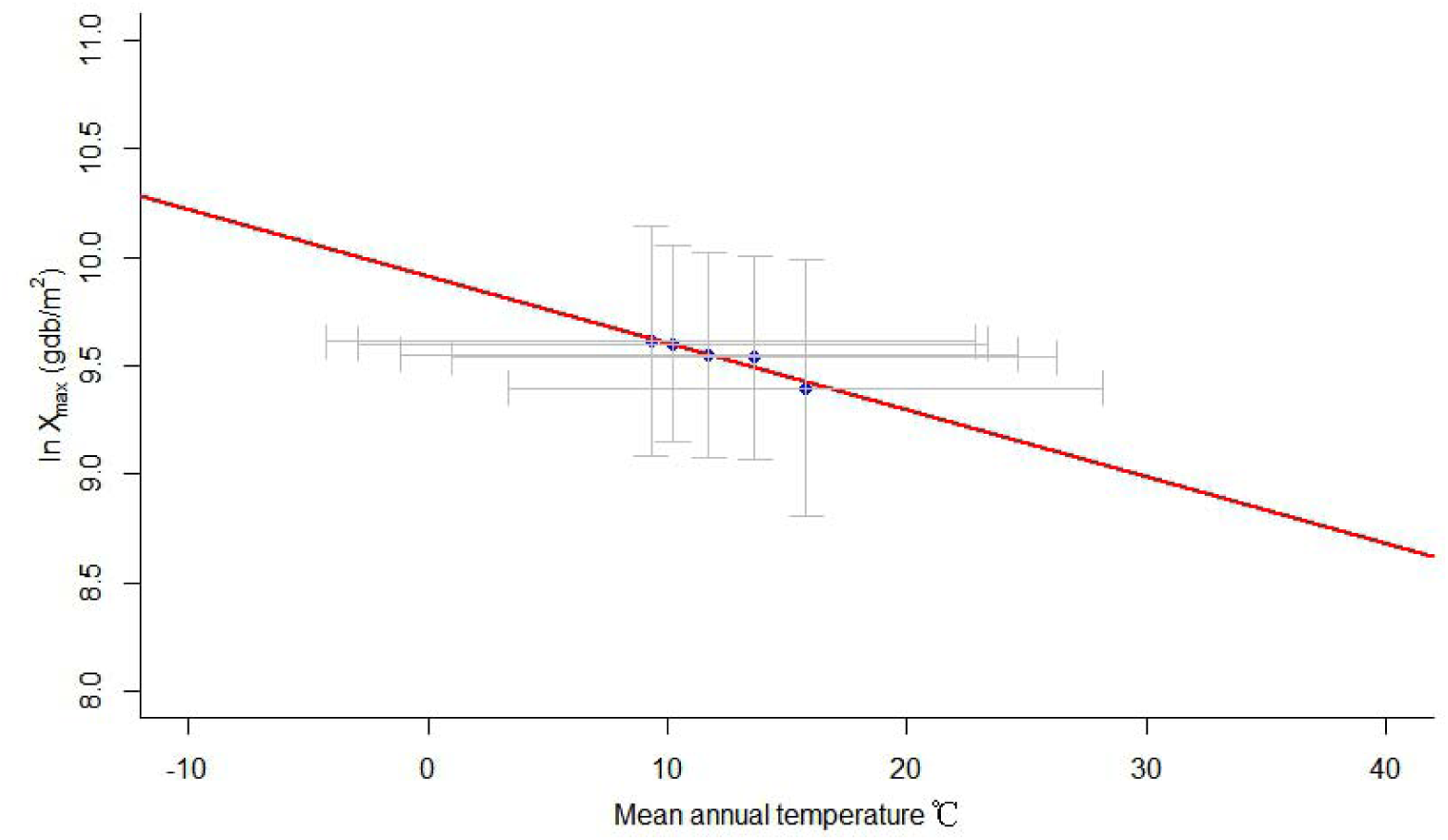
Mean annual temperature and WPC maximum biomass for 20 years from 2020 to 2100. The blue dots represent the means for the 2016–2020, 2020–2040, 2040–2060, 2060– 2080, and 2080–2100 periods from left to right. The gray line represents the standard deviation, and the red line is *y* = −0.029*x* +10.18 (R^2^ = 0.75, *P* = 0.04).

**Fig. S7.**
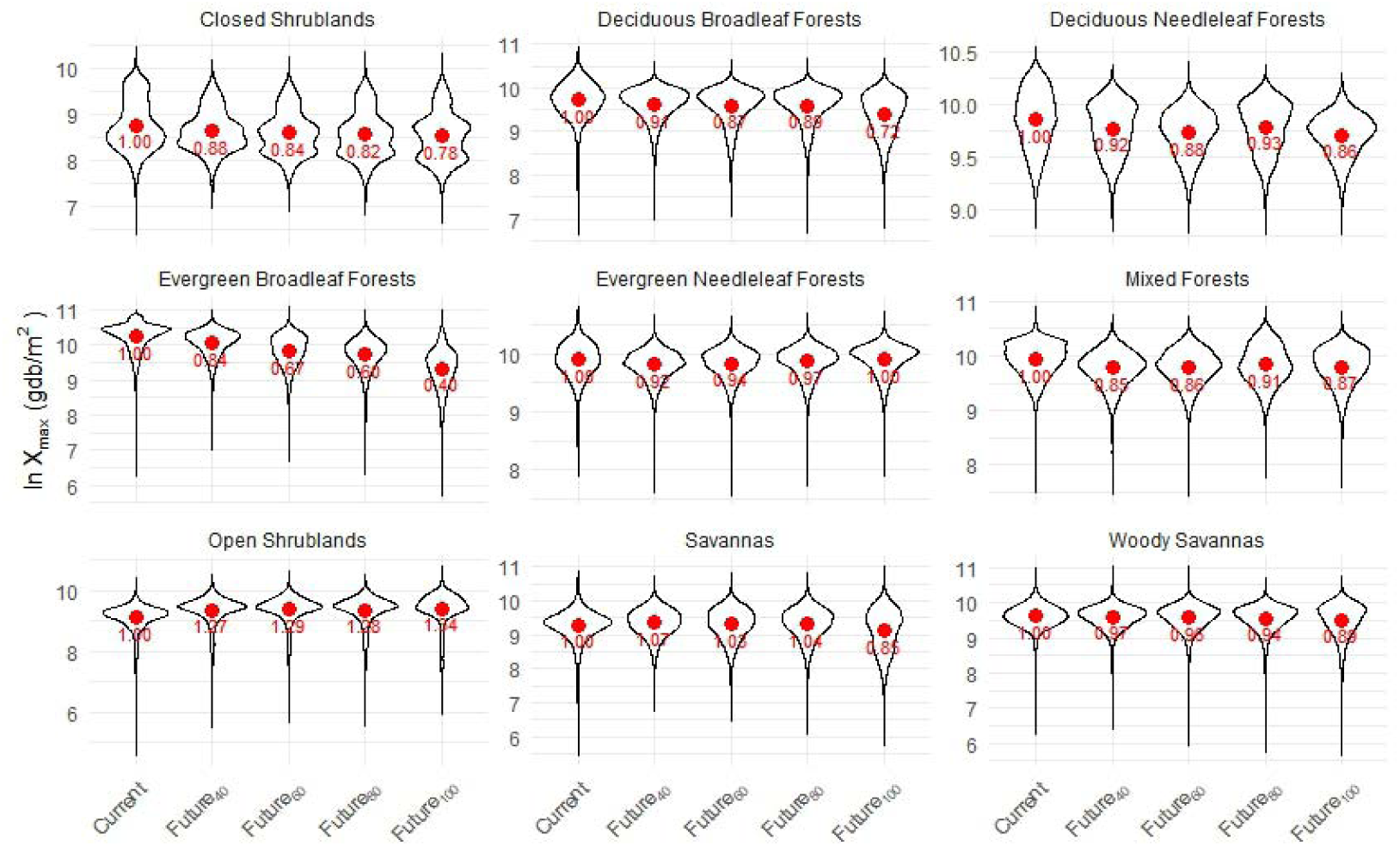
Future changes in the maximum biomass of different WPCs. The red dots indicate the mean values, and the numbers below them represent the ratios of the mean maximum biomass in different periods to the mean maximum biomass in 2020. Future_2040_, Future_2060_, Future_2080_, and Future_2100_ for 2020-2040, 2040-2060, 2030-2080 and 2080-2100, respectively.

**Fig. S8.**
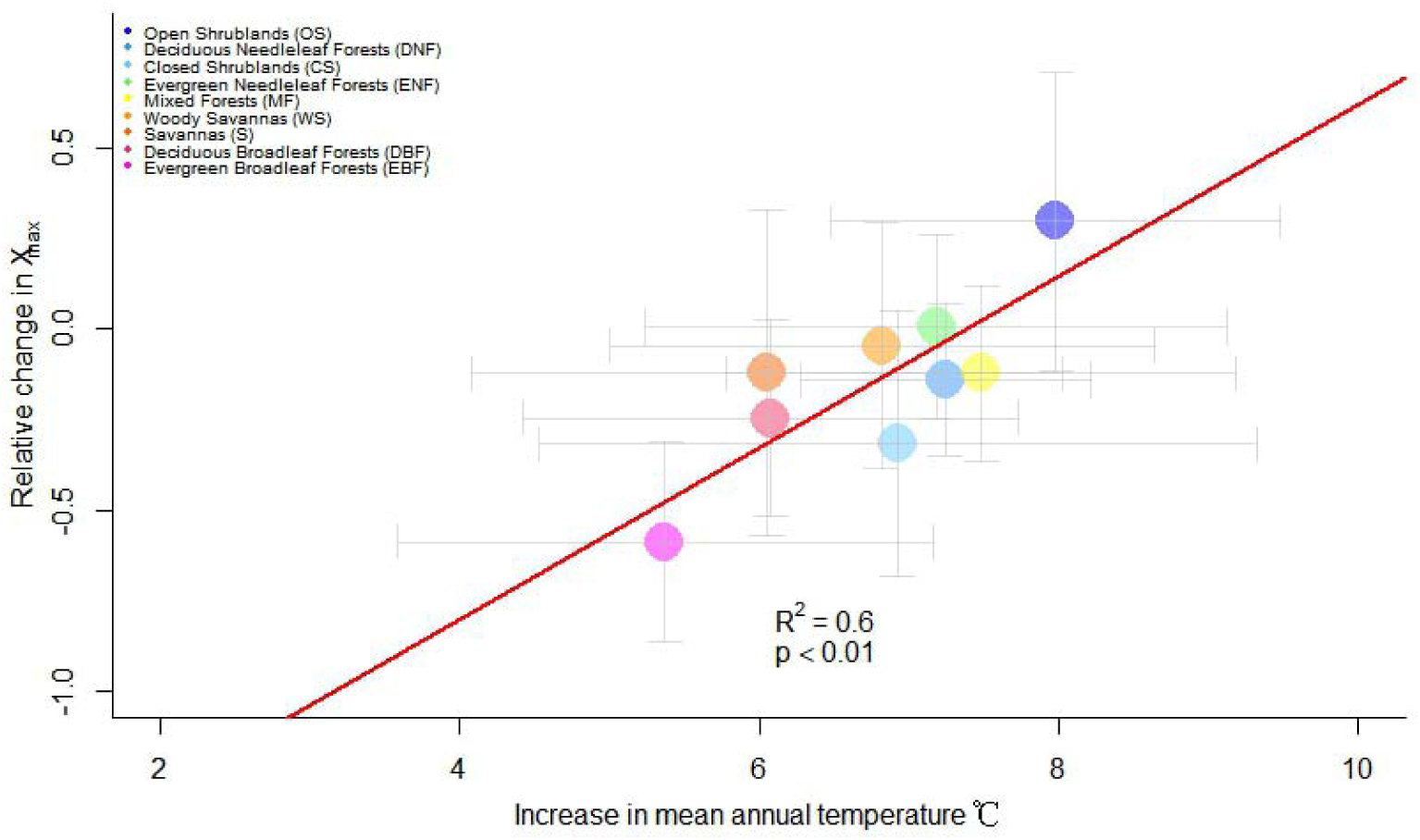
Relative change in *M*_max_ for different WPC types under mean annual temperature increase during 2020 to 2100. The relative change in *M*_max_ is defined as (*M*_max (2100)_-*M*_max_)/*M*_max_, where *M*_max (2100)_ is the mean *M*_max_ from 2080 to 2100. The red solid line is the fitted line. The short grey line is the standard deviation.

**Fig. S9.**
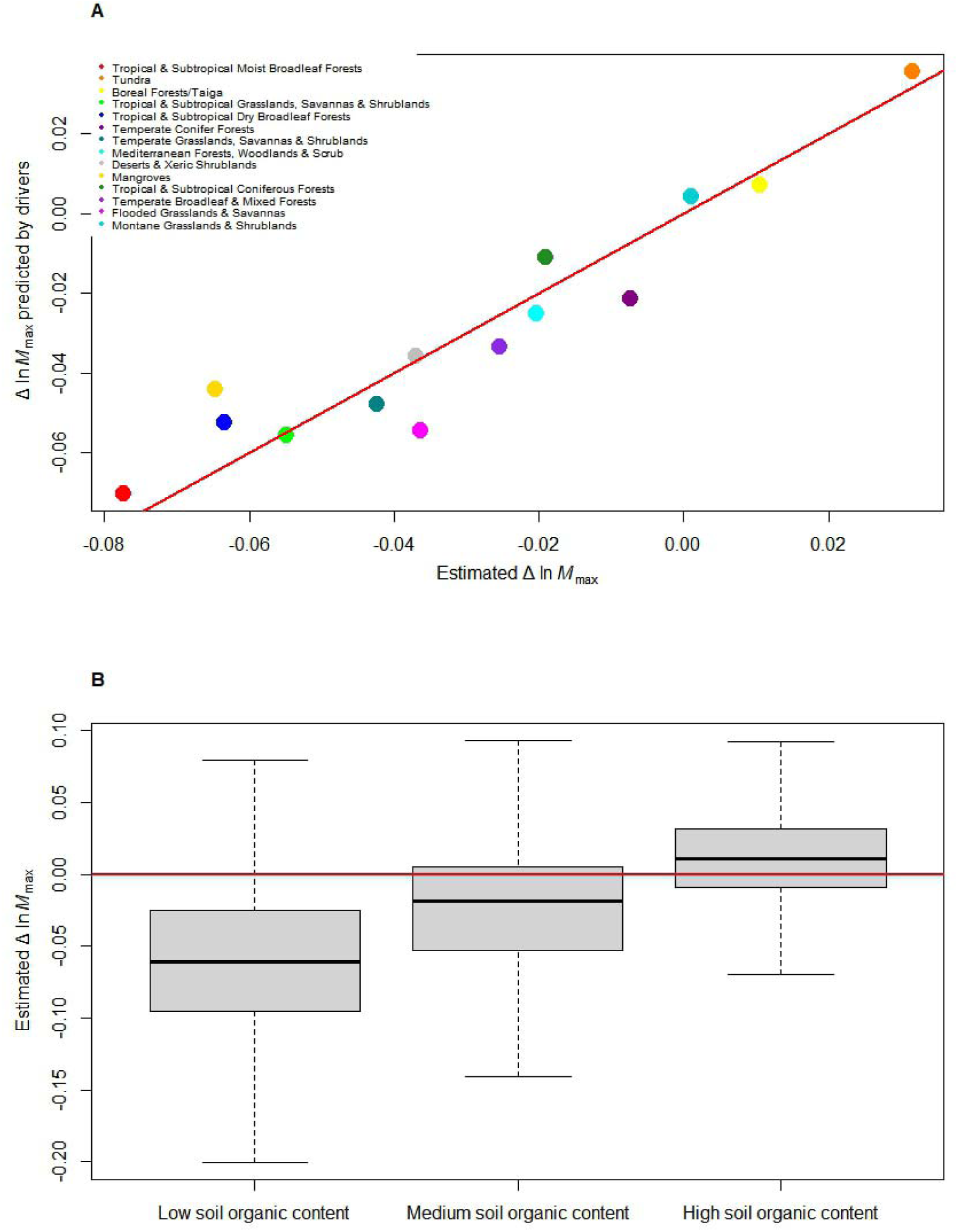
Multiple linear regression explaining _Δ_ln *M*_max_ at the ecoregion scale (A) and across different levels of soil organic matter content (B). A, B: Δln *M*_max_, relative change, defined as ln *M*_max (2100)_/ln *M*_max_ - 1, where *M*_max (2100)_ is the mean *M*_max_ from 2080 to 2100. B:The low, medium, and high classes correspond to soil organic matter content in the quantiles less than 0.25, between 0.25 and 0.75, and above 0.75, respectively.

**Fig. S10.**
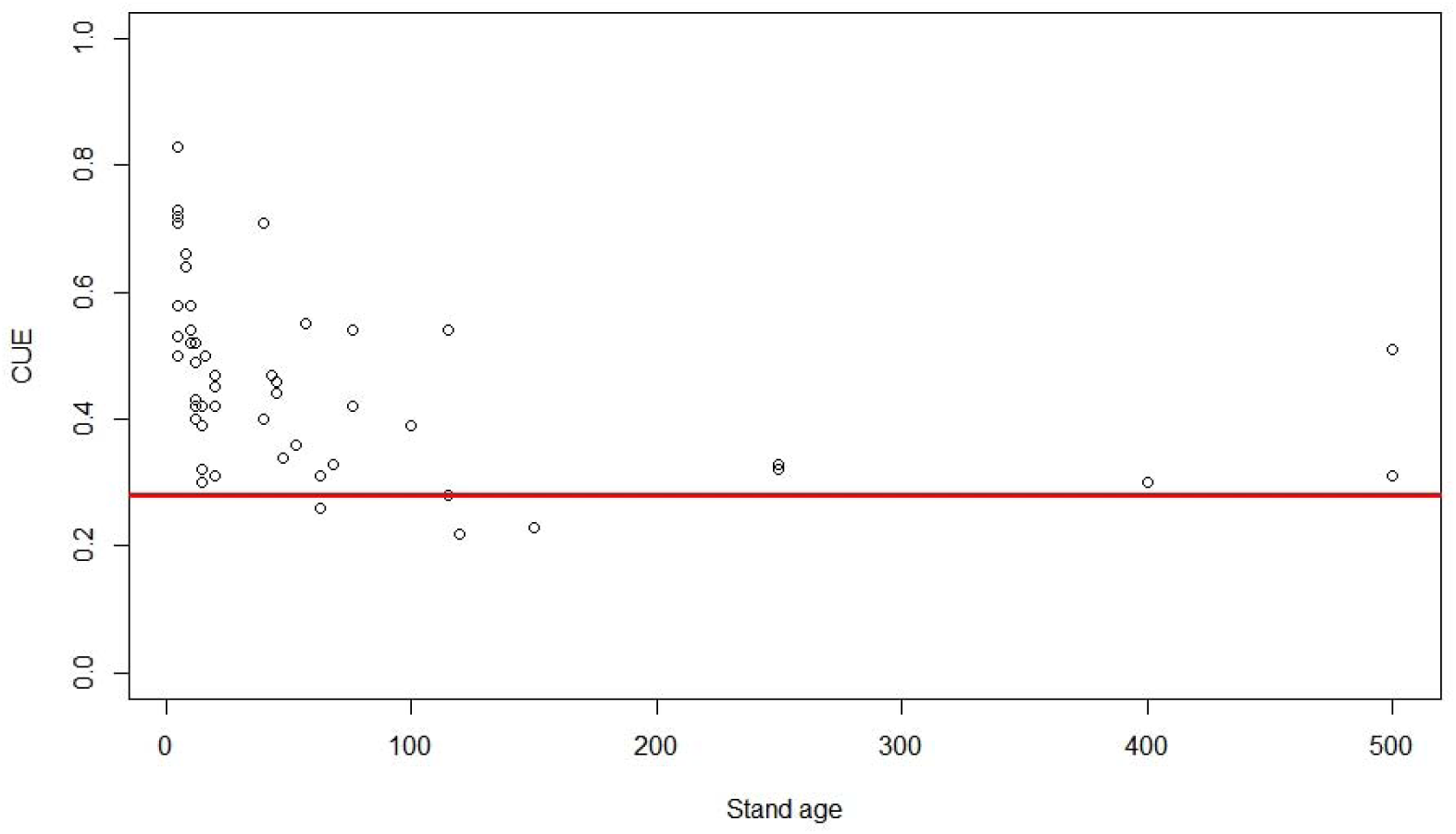
Forest carbon use efficiency vs. stand age. Carbon use efficiency (CUE) is the ratio of NPP to GPP (data from ref. (DeLucia *et al*. 2007) Table 1), and here, NPP is equivalent to our NPP_s._ The red horizontal line is *y* = 0.28

**Fig. S11.**
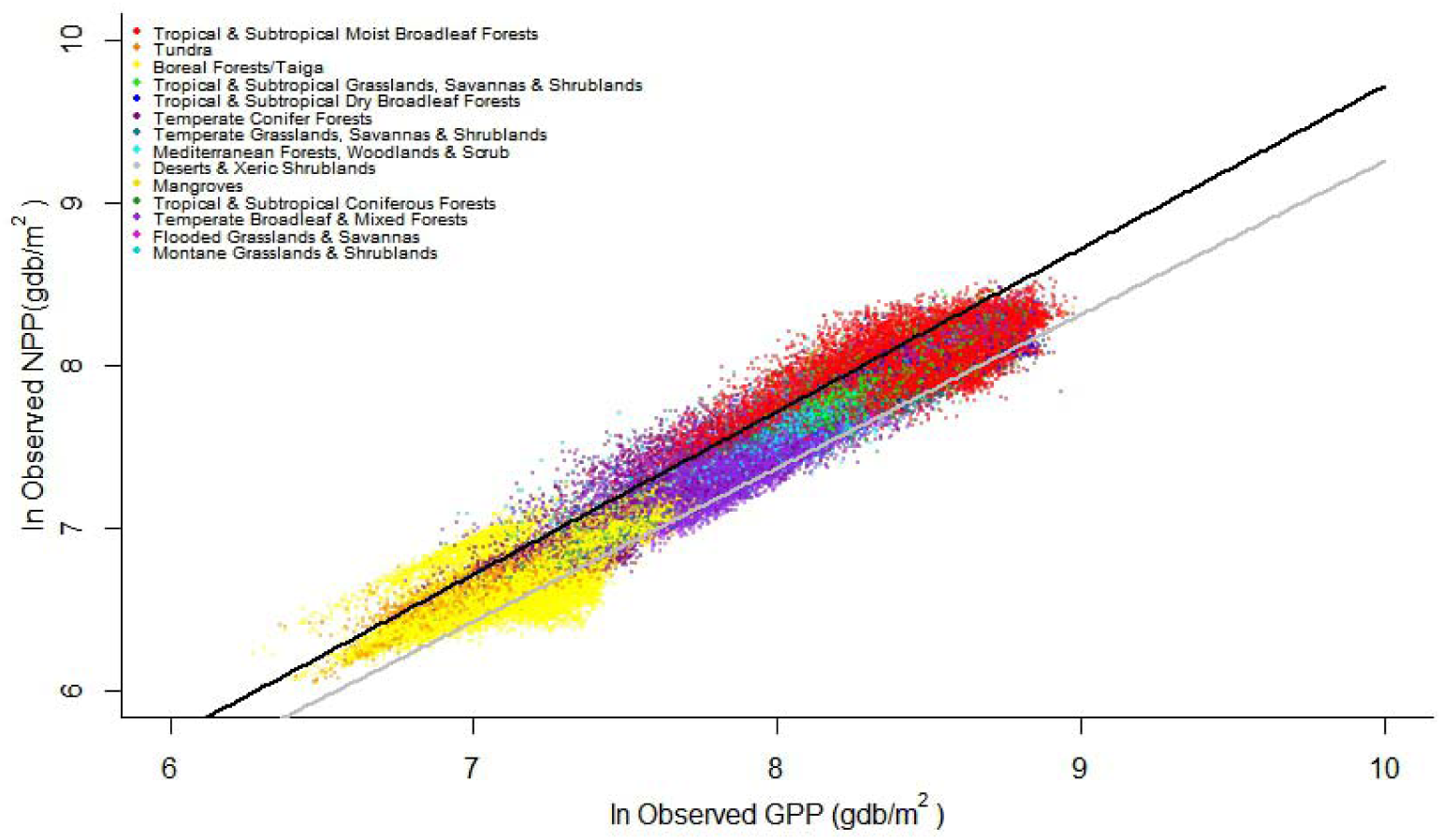
Relationship between NPP raster data close to the NPP predicted by Eq. S6 and their corresponding GPP raster data. The grey and black solid lines represent the 0.95 and 0.05 quartile fit curves. The equations are *y* = *x* - 0.29 and *y* = 0.95*x* - 0.18. In the Cartesian coordinate system, the intercepts of these fitted curves correspond to slopes of 0.75 and 0.84, respectively, which aligns with theoretical predictions (0.64 - 0.83).

